# Anatomy, histology and ultrastructure of the adult human olfactory peduncle: blood vessel and *corpora amylacea* assessment

**DOI:** 10.1101/2024.11.26.625157

**Authors:** JF. Raspeño-García, S. González-Granero, V. Herranz-Pérez, A. Cózar-Cuesta, E. Artacho-Pérula, R. Insausti, JM. García-Verdugo, C. de la Rosa-Prieto

## Abstract

The mammalian olfactory system is responsible for processing volatile chemical stimuli and is composed of several structures including the olfactory epithelium, olfactory bulb, olfactory peduncle (OP), and olfactory cortices. Despite the critical role played by the OP in the transport of information it has remained understudied. In this work, optical, confocal, and electron microscopy were employed to examine the anatomy, histology, and ultrastructure of six human OP specimens (ages 37–84 years). Three concentric layers were identified in coronal sections: the external layer (EL), the axonal layer (AL), and the internal layer (IL). Immunohistochemistry revealed the distribution of neurons and glial cells throughout the OP. Two neuronal morphologies were observed: granule cells and larger pyramidal cells, the latter associated with projection neurons of the anterior olfactory nucleus. Astrocytes were uniformly distributed with a more radial morphology in the EL. Oligodendrocytes were mainly located in the AL. Blood vessels (BVs) were evenly distributed along the OP, with a mean luminal area of 82.9 µm² and a density of 1.26 %, with a significant increase in the IL. CA were abundant, with an average size of 49.3 µm² and a density of 3.23 %. CA clustered near BVs, particularly at tissue edges, with both size and density increasing with age. Notably, CA showed strong associations with astrocytes. This study provides the first detailed qualitative and quantitative data on the internal organization of the human OP, which may contribute to a better understanding of the pathophysiology of some neuropathological disorders.

## 1. Introduction

The olfactory system is the organ responsible for detecting volatile substances in the surrounding environment. In humans, it plays a role in olfactory-associated functions such as odor discrimination, memory, and emotional processing. This system is composed of the olfactory epithelium, the olfactory bulb (OB), the olfactory peduncle (OP) (Fig.1), and the olfactory cortices (Paxinos 2014). From the olfactory epithelium, located in the nasal cavity, sensory olfactory neurons project to the OB through the cribriform plate of the ethmoid bone, forming the primary projections. This olfactory information is transmitted centripetally along the entire length of the OP, also referred to as the olfactory tract, to the primary olfactory cortex, which includes the piriform cortex, entorhinal cortex, periamygdaloid cortex, anterior olfactory nucleus (AON), and olfactory tubercle. The OP serves as the essential conduit for these signals, ensuring their delivery to higher olfactory processing areas. Afterward, the primary olfactory cortex projects to other brain regions such as the orbitofrontal cortex, insular cortex, thalamus, hippocampus, and hypothalamus. Additionally, the OP is not just a route for centripetal connections but also carries centrifugal axons originating from various areas, including the contralateral OB, the horizontal limb of the diagonal band, the amygdala, the hippocampus, the locus coeruleus, and the raphe nuclei. These pathways highlight the bidirectional nature of information flow through the OP. (Saiz-Sanchez D. et al. 2010; Paxinos 2014; Smith y Bhatnagar 2019; Ubeda-Bañon et al. 2020). So that this functional and anatomical arrangement underlines the OP’s crucial role as the main hub connecting various olfactory and related neural systems.

**Figure 1.**
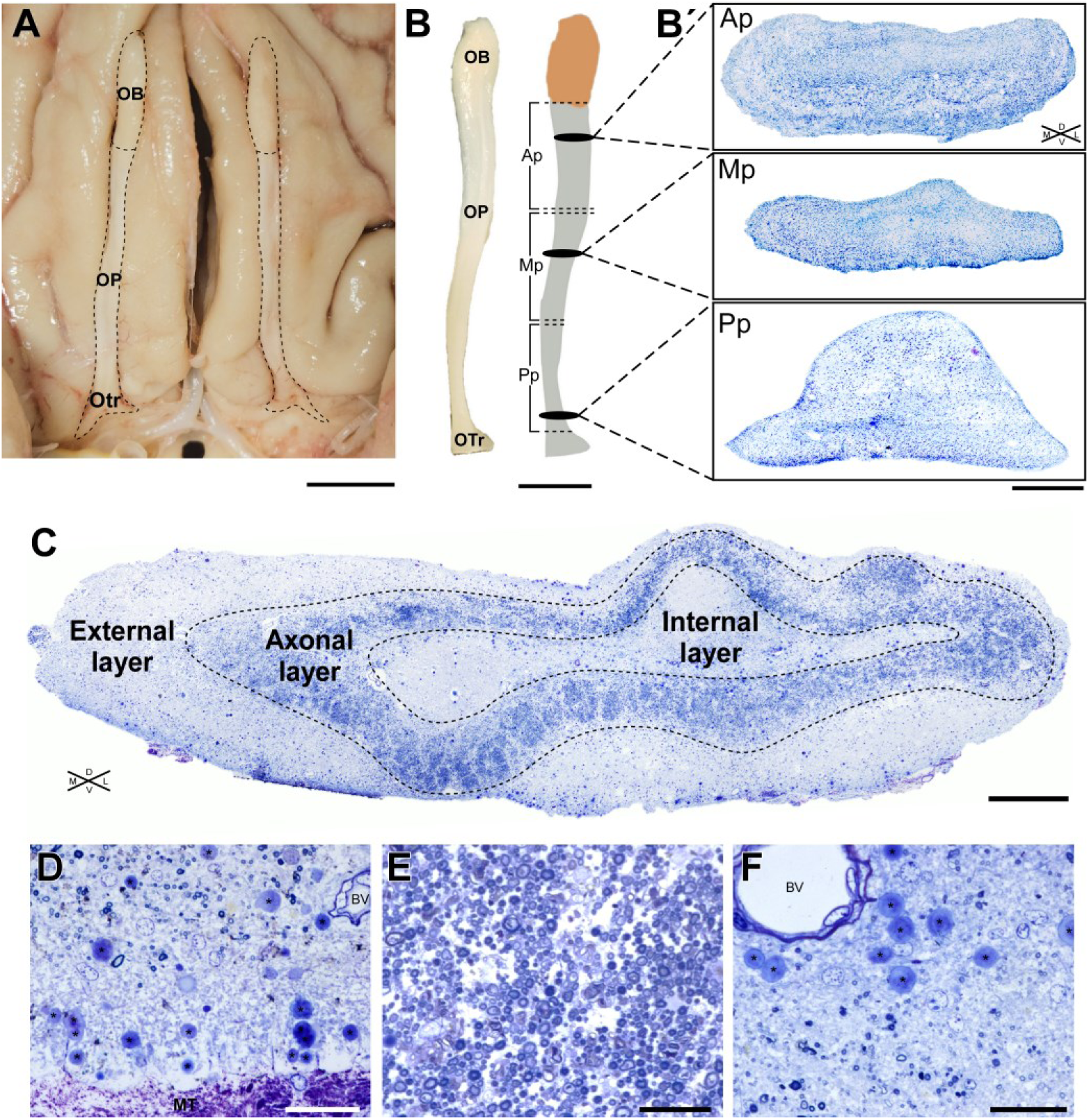
Gross anatomy of the human olfactory nerve (ON). **A)** Inferior view of an adult human brain highlighting the location of the first cranial nerve (CN I), including the olfactory trigone (OTr), olfactory peduncle (OP), and olfactory bulb (OB). **B)** Detailed view of the dissected CN I, showing the three longitudinal portions of the OP: anterior (Ap), intermediate (Mp),and posterior (Pp). **B’)** Representative coronal sections of each portion stained with thionine. **C)** Panoramic micrograph of a coronal semithin section from the anterior portion (Ap), stained with toluidine blue, showing the three histological layers (dashed lines). **D)** External layer (EL) in contact with meningeal tissue (MT), well-vascularized with frequent corpora amylacea (CA), noted by (*) ; small cell bodies dominate. **E)** The axonal layer (AL) contains myelinated axons and numerous small and large cell bodies. **F)** Internal layer (IL) is composed of dense dendritic and axonal processes, featuring the largest blood vessels (BVs) and a high density of CA (*), which frequently cluster around the BVs; small cell bodies are also common. **Abbreviations:** Ap, anterior portion; BV, blood vessel; Mp, intermediate portion; MT, meningeal tissue; OB, olfactory bulb; OC, optic chiasm; OP, olfactory peduncle; Otr, olfactory trigone Pp, posterior portion. **Section thickness:** B’ and C-F) 1.5 µm. Scale bars: A and B) 1 cm; B’ 500 µm; C) 200 µm; D-F) 50 µm.

In this regard, the AON is an intriguing structure embedded within the OP. The AON is part of the anterior olfactory cortex and acts as a laminated structure that connects olfactory processing centres across both hemispheres. Furthermore, the AON is subdivided into several regions, each with distinct architectures and connectivity patterns. These subregions undoubtedly contribute to the modulation of olfactory inputs, facilitating communication between the OBs and enhancing bilateral olfactory integration and processing (Berendsen et al. 2015; Brunert, Medinaceli Quintela, y Rothermel 2023).

To date, beyond the AON, only a few neuroanatomical investigations have been conducted strictly on the olfactory OP components. It is notable that studies which do mention the OP often do so in the context of broader olfactory pathways, rather than in the form of detailed anatomical investigations. We present here some anatomical and cytomorphological features that will help us better understand this structure, which has traditionally been considered merely a passage zone for the olfactory tract.

### 1.1. Blood supply

In regard to the vascular supply of the OB and OP, previous studies indicated that the olfactory epithelium and ventral aspect of the OB receive a vital extracranial blood supply from the sphenopalatine artery, in addition to the anterior and posterior ethmoidal arteries (Bartholdy 1897; Osborn 1978). In the intracranial space, the olfactory artery anastomoses with the OP and OB via the arachnoid membrane, thereby supplying the entire structure (Favre et al. 1995; Hendrix et al. 2014). However, there is a considerable lack of information on the internal distribution of blood flow in the OP. In this study we present, for the first time in humans, a quantification of the BVs and their internal distribution.

### 1.2. Corpora amylacea

CA are anucleate fibrous rounded or ovoid inclusions that present a wide range of sizes, which appear to be related to the tissue in which they accumulate. They have been observed with diameters ranging from 2 to 20 µm in the retina (Avendano et al. 1980) to 80 to 160 µm in the lungs (Martínez-Girón y Pantanowitz 2021). Furthermore, numerous studies have confirmed the presence of CA in the central nervous system. They are frequently observed in the hippocampus, subependymal regions of the ventricles (Alder 1953; Sakai et al. 1969b; 1969a) as well as in the glia limitans in close proximity to the pia mater (Sbarbati et al. 1996); proximity to BV and the glymphatic system, primarily within the cytoplasm of perivascular glial cells (Riba, Del Valle, et al. 2022). Additionally, they have been observed in the olfactory tract (Bathini et al. 2019) or the brainstem, specifically the *substantia nigra pars compacta* and the *canalis centralis.* (Navarro et al. 2018). Some of these studies evaluate the average diameter of the identified CA. For example, Xu et al. (Xu et al. 2021) measured the diameter of CA in the hippocampal formation at 8 to10 µm, while Bathini et al. (Bathini et al. 2019) measured the diameter in the OP at 14 to 16 µm. Despite the lack of consensus regarding their origin and function, there is a general agreement that they may play a significant role in the accumulation of cellular metabolic products or other substances present in the microenvironment (Cavanagh 1999; Augé et al. 2018). Other authors have proposed that they may be formed as a result of cellular or axonal degradation (Anzil et al. 1974; Song et al. 2014). It is also thought that CA accumulate in tissues as part of the normal aging process (Mrak, Griffin, y Graham 1997). However, numerous studies have also linked their presence in specific locations to pathological conditions (Riba et al. 2021), or noted differences in their number, size, or elemental composition compared to control populations. CA have also been associated with various neuropathological conditions including Alzheimeŕs disease (Singhrao, Neal, y Newman 1993; Renkawek y Bosman 1995; Wander et al. 2022), Parkinsońs disease (Mizutani, Satoh, y Morimatsu 1987; Buervenich, Olson, y Galter 2001) amyotrophic lateral sclerosis (Atsumi 1981; Gáti y Leel-Ossy 2001), and others.

Similar to the BVs, we present a quantitative approach to the distribution, size, and density occupied by the CA along the OP.

### 1.3. Olfactory peduncle anatomy and histology

The anatomy and histology of the OP have been insufficiently studied, as commented. There are few studies, to the best of our knowledge, that have focused on describing the histological and anatomical organization of the OP in humans (Crosby y Humphrey 1941; Paxinos 2014). Furthermore, there is a lack of research on how this could vary in both normal aging processes and pathological situations.

This study presents a detailed anatomical, cytoarchitectural, and ultrastructural analysis of the adult human OP along its rostrocaudal axis, from the caudal limit of the OB to the rostral part of the olfactory trigone. For this purpose, and due to its great length, the OP has been divided into three portions of 1 cm each (approx.): anterior (Ap), intermediate (Mp), and posterior (Pp) portions (Fig. 1). We have investigated the morphology along the rostrocaudal axis, as well as the distribution, cytoarchitecture, and ultrastructure of the primary components, including astrocytes, microglia, oligodendrocytes and neurons, using coronal and longitudinal sections of each portion. Furthermore, we have examined the distribution, number, size, density, and ultrastructure of the BV and CA.

## 2. Materials and methods

### 2.1. Tissue

A total of 6 human olfactory peduncles (N= 6), left or right, from individuals of both sexes (3 male, 3 female) ranging from 37 to 84 years old and undiagnosed with neurological pathologies, were used in this study (Table 1A). The tissue was obtained from archives and the donation program established at the Faculty of Medicine in Albacete, University of Castilla-La Mancha; with the appropriate informed consent from the families prior to sample donation. All procedures were approved by the local Committee on the Ethics of Medicinal Products Research (CEIm).

**Table 1.**
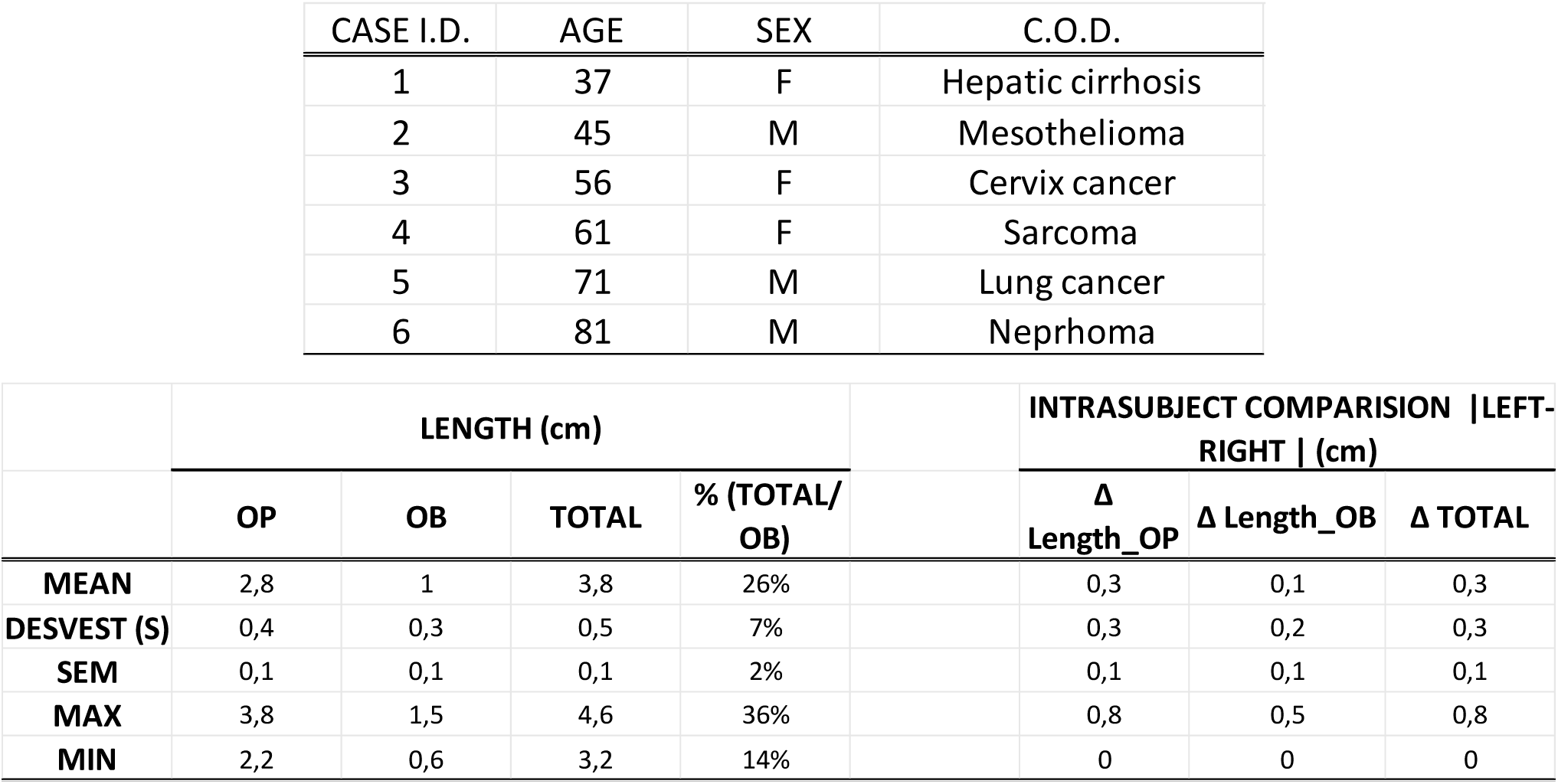
Population and olfactory nerves measures. **A)** Population sample used in the study. **B)** Summary of the measurements obtained for each olfactory peduncle (OP) and bulb (OB), including the total length of the structure, the length of the peduncles, the length of the bulbs, and the percentage contribution of the bulb to the total length of the structure, measured from the most rostral part of the olfactory trigone. Additionally, a comparison of the morphological variation typically observed between the lengths of the left and right organs within the same individual is provided.

### 2.2. Perfusion and fixation

We processed the cadaveric tissue using a previously published protocol (Insausti et al. 2023), which allows differential perfusion of brain structures with respect to the rest of the body. This procedure was performed within a short post-mortem interval of 4-8 hours, which ensures proper tissue fixation, enhanced ultrastructural preservation, and allows for tissue homogeneity in comparisons.

### 2.3. Optical and confocal microscopy

For optical and confocal microscopy, the tissue was cryoprotected in 0.1 M phosphate-buffered (PB) solution with 30% sucrose at 4°C for 24-48 hours. Subsequently, the tissue was sectioned at 30 µm using a freezing microtome and collected in 24-well plates with LRC solution (30% v/v ethylene glycol, 20% v/v glycerol, 0.05 M PBS, pH 7.4) and then stored at 4°C. Some sections were counterstained with thionine for morphology studies. Coronal semithin sections were used for cytoarchitecture analysis and assessment of BVs and *corpora amylacea*. Images were obtained using a *Zeiss Axio Imager M.2 with APOTOME* and composed with *Zeiss Tiles* software.

### 2.4. Immunofluorescence detection

For immunofluorescence analysis, 30-micrometer-thick sections from 6 olfactory peduncles, previously mounted on gelatinized glass slides, were utilized. The sections were subjected to 6 washes of 5 minutes each in 0.1 M phosphate-buffered saline (PBS), pH 7.4, at room temperature. Subsequently, 6 washes were performed in PBST (0.1 M PBS pH 7.4, 0.1% Triton) for 5 minutes each at room temperature. Thereafter, the tissue sections were encircled with a hydrophobic pen and incubated for 1 hour at room temperature, covered with a blocking solution of PBST, 1% bovine serum albumin (BSA), and 10% normal horse serum (NHS). After removing the blocking solution, the primary antiserum, consisting of primary antibodies added at the established dilutions in PBST with 2% NHS, was incubated at 4°C for the time established for each antibody (Table 2A) with gentle agitation. Following incubation with the primary antibody, the sections were washed 6 times for 5 minutes in PBST under agitation. Subsequently, the corresponding secondary antiserum (Table 2B) was added and incubated in the dark for 1.5 hours at room temperature with gentle agitation. The sections were then washed in 0.1 M phosphate-buffered saline (PBS) to remove the antiserum. Finally, the sections were incubated with 1 µg/mL 4’,6-diamidino-2-phenylindole (DAPI) in 0.1 M PBS for 1 minute and rinsed in 0.1 M PB, pH 7.4 for 5 minutes. To observe and preserve the slides, we utilized PVA-DABCO (polyvinyl alcohol mounting medium and 1,4-diazabicyclo [2.2.2] octane) as the mounting medium.

**Table 2.**
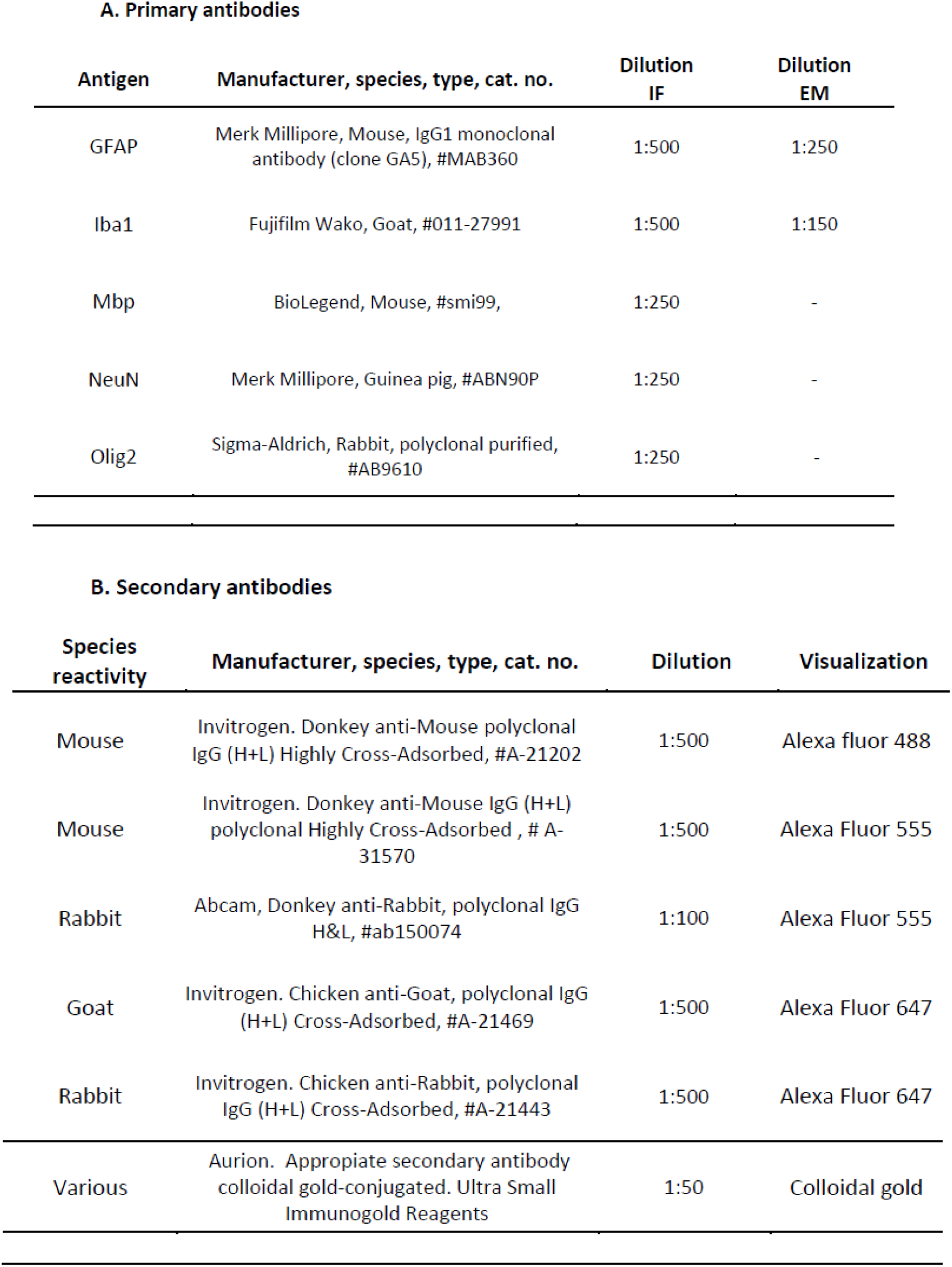
Primary A), and secondary B) antibodies.

A. Primary antibodies
| Antigen | Manufacturer, species, type, cat. no. | Dilution<br>IF | Dilution<br>EM |
| --- | --- | --- | --- |
| GFAP | Merk Millipore, Mouse, IgG1 monoclonal antibody (clone GA5), #MAB360 | 1:500 | 1:250 |
| Iba1 | Fujifilm Wako, Goat, #011-27991 | 1:500 | 1:150 |
| Mbp | BioLegend, Mouse, #smi99, | 1:250 | - |
| NeuN | Merk Millipore, Guinea pig, #ABN90P | 1:250 | - |
| Olig2 | Sigma-Aldrich, Rabbit, polyclonal purified, #AB9610 | 1:250 | - |

Table 2. – Primary A), and secondary B) antibodies.
| Species reactivity | Manufacturer, species, type, cat. no. | Dilution | Visualization |
| --- | --- | --- | --- |
| Mouse | Invitrogen. Donkey anti-Mouse polyclonal IgG (H+L) Highly Cross-Adsorbed, #A-21202 | 1:500 | Alexa fluor 488 |
| Mouse | Invitrogen. Donkey anti-Mouse IgG (H+L) polyclonal Highly Cross-Adsorbed , # A-31570 | 1:500 | Alexa Fluor 555 |
| Rabbit | Abcam, Donkey anti-Rabbit, polyclonal IgG H&L, #ab150074 | 1:100 | Alexa Fluor 555 |
| Goat | Invitrogen. Chicken anti-Goat, polyclonal IgG (H+L) Cross-Adsorbed, #A-21469 | 1:500 | Alexa Fluor 647 |
| Rabbit | Invitrogen. Chicken anti-Rabbit, polyclonal IgG (H+L) Cross-Adsorbed, #A-21443 | 1:500 | Alexa Fluor 647 |
| Various | Aurion. Appropriate secondary antibody colloidal gold-conjugated. Ultra Small Immunogold Reagents | 1:50 | Colloidal gold |

### 2.5. Electron microscopy

Ultrastructural studies were conducted, and for this purpose, the tissue was post-fixed in 2% paraformaldehyde and 2.5% glutaraldehyde in 0.1 M PB for 7 days. Subsequently, the tissue was washed with 0.1 M PB, post-fixed with 2% osmium tetroxide, rinsed, dehydrated, and embedded in Durcupan epoxy resin (Sigma). Semithin sections (1.5 µm) were obtained with a diamond knife on an ultramicrotome (Leica UC7), stained with 1% toluidine blue, and examined under an optical microscope (Nikon E-800). Later, ultra-thin sections (60–70 nm) were obtained, stained with lead citrate (Reynolds solution), and examined under a transmission electron microscope (*FEI Tecnai G^2^ Spirit BioTwin*). Images were acquired using Radius software (Version 2.1) with an XAROSA digital camera (EMSIS GmbH, Münster, Germany). For pre-embedding immunohistochemistry, tissue was fixed with 4% PFA. Pre-embedding immunogold staining was performed by incubating sections in primary antibodies (Table 2A) and the appropriate colloidal gold-conjugated secondary antibodies (1:50; UltraSmall; Aurion) (Table 2B) as previously described (Sirerol-Piquer et al. 2012).

### 2.6. Image analysis

For quantitative analysis (N_cases_=6; n_sections_=18) of BVs and CA, each OP was represented by three semithin coronal sections (1.5 µm) from the anterior (Ap), intermediate (Mp), and posterior (Pp) portions (Fig. 1B,B’). After staining with toluidine blue, panoramic images of the OP were captured using a *Zeiss Axio Imager M.2* with *APOTOME* and *ZEN* software. Subsequently, the sections were analyzed for coronal area and layer delineation. The different OP layers (s*upp. mat. 1*), BVs, and CA were identified and outlined with *Canvas X Pro software*. The vectorial polygons were assessed using an *ImageJ (DOI:10.5281/zenodo.13969287)*.

This methodology allowed detailed enumeration and assessment of the dimensions and positioning of each element within the layers. Data recorded included case, OP portion, layer, count (F), individual area (A), and density (D), which measures the percentage of tissue occupied by BVs (*supp. mat. 2)* or CA (*supp. mat. 3)* relative to the total section or layer (*supp. mat. 4)*. Results has been stored in a public repository (*DOI:10.5281/zenodo.13969287*).

### 2.7. Statistical analysis

Data analysis was performed using GraphPad Prism 20.00. Descriptive statistics included mean, median, standard error (SEM), standard deviation (SD), sample variance, skewness, and range. Normality was assessed with the D’Agostino & Pearson omnibus normality test. Group comparisons used Kruskal-Wallis (KW) and Dunn’s test (DT) or one-way ANOVA and Tukey’s test (Tk HSD), depending on normality (*supp. mat. 5 and 6)*. Results are shown as mean ± SEM. Pearson’s test (PT) was used for linear correlations with normality, and Spearman’s test (ST) was used when normality assumptions were not met (*supp. mat. 7)*.

## 3. Results

### 3.1. Morphology of the human adult OP

The entire structure, including the OP and the OB, is located in the olfactory sulcus surrounded by meningeal tissue, pia mater and arachnoid (Fig. 1A, D). From an external perspective, a dorsal protuberance is apparent along the rostrocaudal axis of the OP, particularly in the intermediate and posterior portions (Fig. 1B’, C), which conforms to the space of the olfactory sulcus. This morphology gradually diminishes as we approach the terminal region of both the olfactory trigone and the OB, which aids in the orientation of coronal slices. The OP exhibits a flattened dorsoventral morphology in intermediate areas and a rounded morphology towards the ends (*supp. mat. 8*). The OP and OB differ in length (*supp. mat. 9)* between individuals and even between the left and right sides of the same individual (Table 1B). Following the dissection of each pair of nerves from each subject (Table 1B) at the level of the olfactory trigone, the length of the OP and OB was assessed, with values ranging from: 3.2 to 4.6 cm for the entire structure (MEAN_total_=3.8±0.1 cm); 2.2-3.8 cm (MEAN_OP_=2.8±0.1 cm) the OP; and 0.6-1.5 cm (MEAN_OB_=1.0±0.1 cm) the OB (see Table 1B). A comparison of the total structure length reveals that the proportion represented by the OP and the OB also varies. As observed, the length of the OB exhibits greater variability between cases compared to the OP, with coefficients of variation (CV) of 30% and 14%, respectively.

### 3.2. Cytoarchitecture

The OP has been studied under optical, confocal, and electron microscopy on coronal sections, and three different layers have been identified: external (EL), axonal-intermediate (AL), and internal (IL) layer. These layers, arranged concentrically, show variations in thickness along the OP. In this study, we have been measured the coronal area of each section, layer areas, and their proportions regarding to the total area of each section.

Coronal area of OP sections, excluding surrounding meningeal tissue (MT), ranged from 1 to 3 mm² (Fig. 2A), averaging 2.09±0.16 mm² (N=6, n=18; Table 1A). The smallest section was 1.01 mm² and the largest was 3.37 mm². Although the OP sometimes appears narrower in the medial portion, our sample showed no significant differences along its rostrocaudal axis. The distinctive cytoarchitecture and structural composition of each layer are detailed hereafter.

**Figure 2.**
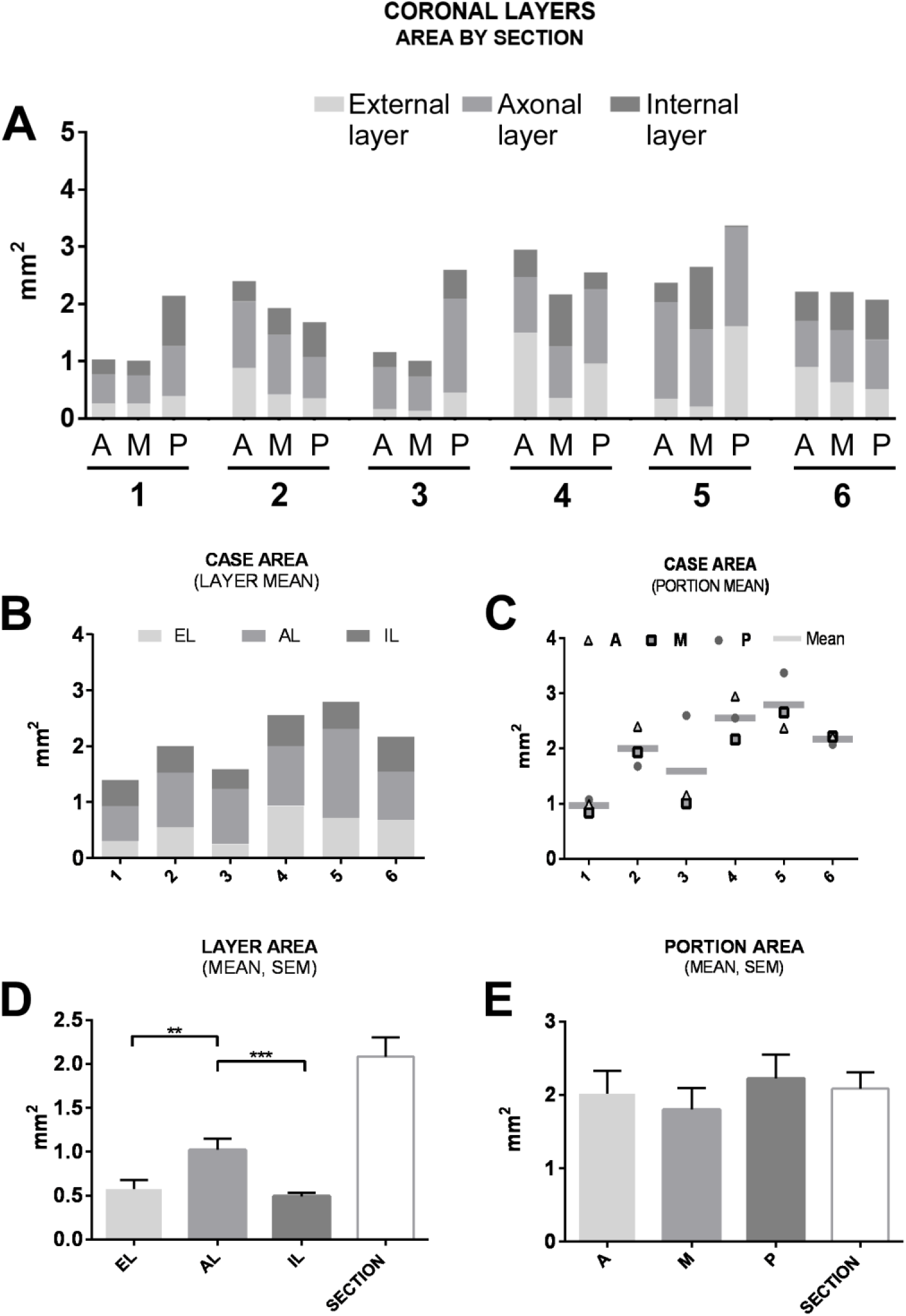
Coronal areas of the olfactory peduncle (OP) across different layers and longitudinal portions. **A)** Area values of each layer (EL, AL and IL) were measured in the three longitudinal portions of the OP (Ap, Mp and Pp) from six OPs (left or right) in six individuals. Each case included three sections (N=6, n=18) corresponding to the longitudinal portions of the OP in one hemisphere. **B)** The total area for each layer was stacked across the three longitudinal portions for each case. Average area values for each layer (EL, AL, IL) were superimposed both across (A) the sections and (B) within each case. **C)** Case values represent the average coronal area of the OP alongside the values for each longitudinal portion. **D)** The axonal layer (AL) is the largest, accounting for roughly half of the total coronal area in the sections analyzed (Kruskal-Wallis test, P = 0.0002; Dunn’s Test). **E)** The mean coronal area of the sections along the rostrocaudal axis (Ap, Mp, Pp) showed no significant differences between sections. **Abbreviations**: EL, external layer; AL, axonal layer; IL, internal layer; Ap, anterior portion; Mp, intermediate portion; Pp, posterior portion.

#### 3.2.1. External layer (EL)

This layer represents the outermost layer of the analyzed sections. An average area of 0.57±0.10 mm^2^ was obtained (Fig. 2D). On average, this EL contributes 25.6% to the total coronal area. In the intermediate portion (Mp) of the OP, both the area and the fraction of the total area appear to be somewhat smaller. This layer extends from the glial-meningeal boundary (MT), the edge of the tissue, to the layer of myelinated axons (Figs. 1C, E; 4E; 8A). Some isolated neurons and myelinated axons are present, and in certain sections, clusters of myelinated axons and neurons appear embedded within stroma (Figs. 3G-L). It is composed mainly of astrocytes (Fig. 4A, C), which show GAP and adherens junctions (Fig. 5E). Microglia (Iba1+ cells) were also observed (Fig. 4B, D; Fig. 6). In terms of vascularization in this layer, the mean number of BVs per section is similar to that of layer IL, and both of these layers show significantly lower values than the counts in AL (Fig. 9C). The lumen area of the vessels was also measured (Fig. 9E). However, the mean lumen area of BV in this layer is not significantly different from that obtained of the other two layers. The density is the lowest among the three layers, slightly lower than in the AL, but exhibits a significant difference compared to the IL, which has the highest BV density (Fig. 9G). The presence of CA is easy to detect here, as they tend to accumulate near the outer boundary of the tissue where the OP glia limitans is imbricated with meningeal tissue (Fig. 1D).

**Figure 3.**
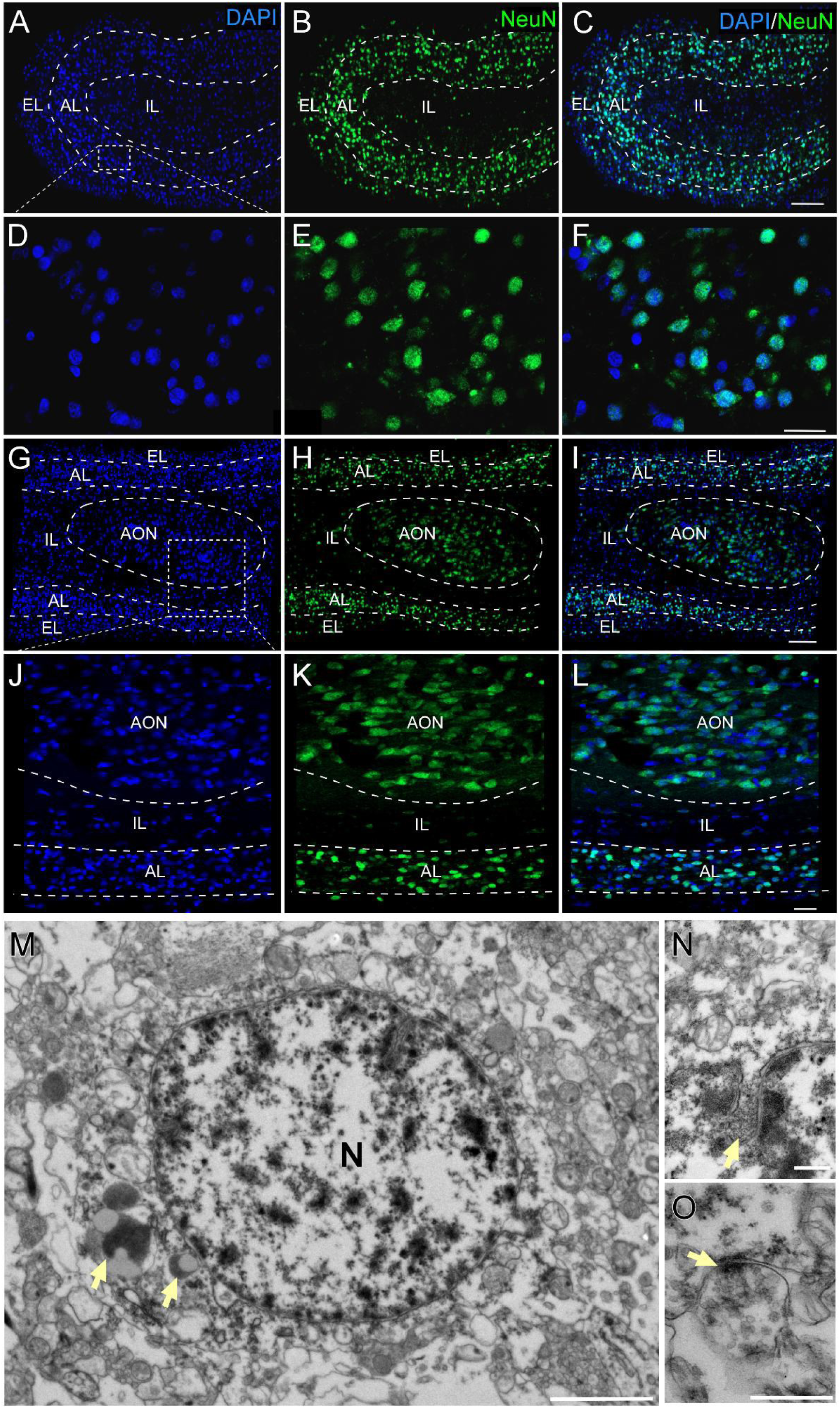
Distribution of the neuronal population in the olfactory peduncle (OP). **A-L)** Immunofluorescence confocal 20x tile composition images. Dotted white squares (A, G) indicate magnified areas (40x), shown in the bottom rows (D-F, J-L). **Coronal plane (A-F):** NeuN-positive cells are mainly located in the axonal layer (AL), corresponding to the distribution of myelinated axons, a pattern consistent throughout the OP. **Parasagittal plane (G-L):** Clusters of NeuN-positive cells (G-I) occasionally appear in the internal layer (IL). These cells (J-L) are larger and pyramidal in shape, differing from the smaller and more abundant granular cells found throughout the OP. These large cells may be part of relay groups in the anterior olfactory nucleus (AON). **M-O)** Transmission electron microscopy images. **M)** Ultrastructure of a typical neuron in the human OP, showing a large cytoplasm with abundant organelles. Lipofuscin bodies (arrows) are frequently seen in adult samples. **N)** Single nuclear invaginations are commonly observed in OP neurons (arrow). **O)** Axo-somatic synapse (arrow), confirming the neuronal identity of this cell type. **Abbreviations:** EL, external layer; AL, axonal layer; IL, internal layer; AON, anterior olfactory nucleus; N, neuron; OP, olfactory peduncle. **Scale bars:** C, I) 100 µm; F, L) 20 µm; M) 2 µm; N, O) 500 nm.

**Figure 4.**
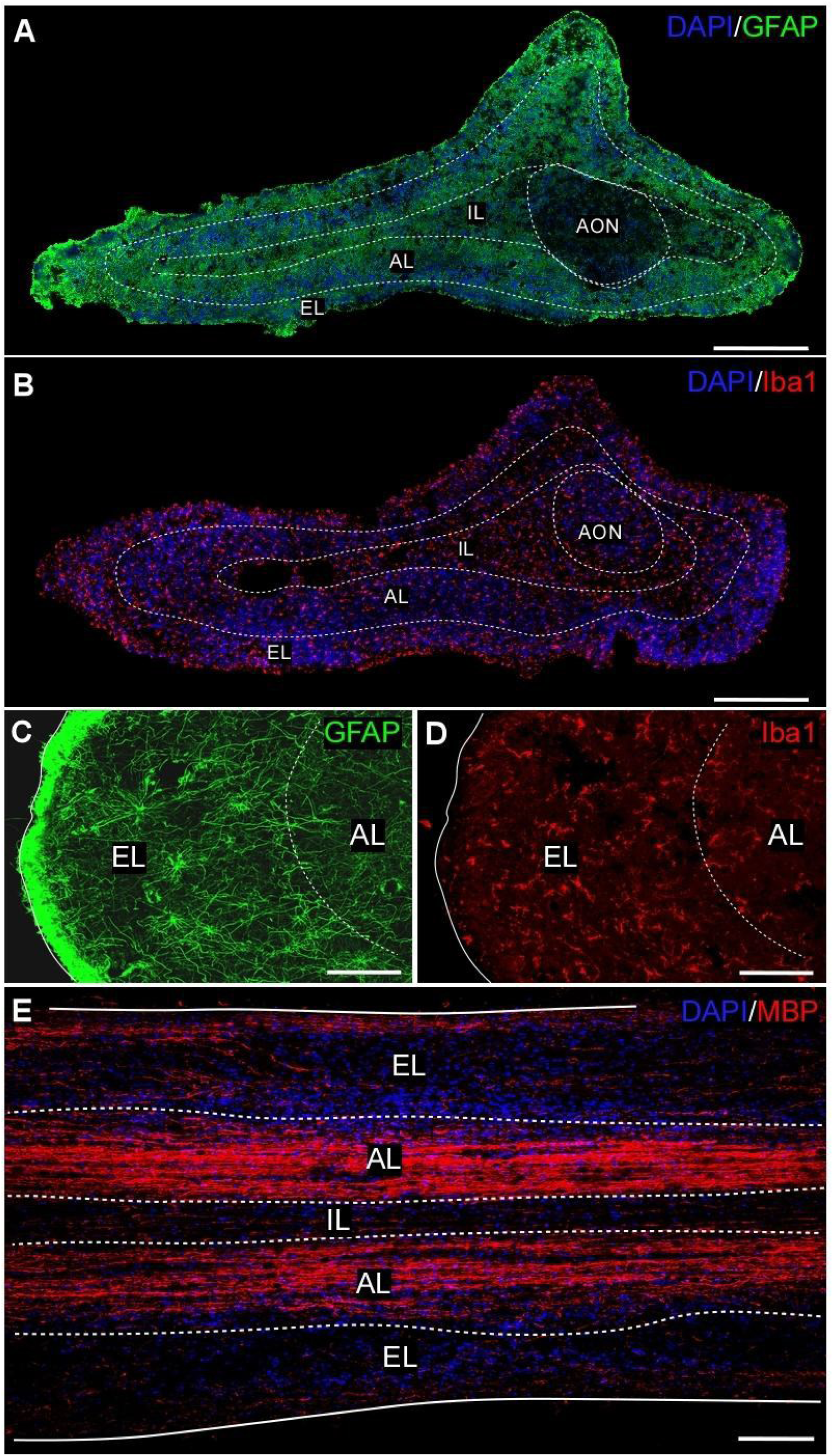
Astrocytic, microglial population, and olfactory tract in the olfactory peduncle (OP). **A)** Confocal image of a coronal section of the OP showing the distribution of GFAP+ cells across the defined layers. A relatively uniform distribution is seen in the three layers (EL, AL and IL), along with the longitudinal axis of the OB. Denser fiber compaction is noted at the tissue periphery (glia limitans), with areas of lower density and a looser glial arrangement observed in some sections, possibly corresponding to portions of the anterior olfactory nucleus (AON). **B)** Distribution of Iba1+ cells, which are evenly spread throughout all regions of the OP, including AON areas. **C, D)** Confocal images combining GFAP and Iba1 markers, illustrating their co-distribution. **E)** Confocal image of a longitudinal section showing myelin fibers positive for MBP. These fibers are mainly organized in two bands forming the axonal layers, with some fibers extending into both the internal and external layers toward the pial tissue. The fiber arrangement is not exclusively longitudinal, as many fibers appear obliquely oriented. **Abbreviations:** AON, anterior olfactory nucleus; AL, axonal layer; EL, external layer; IL, internal layer. **Section thickness:** 30 µm. **Scale bars:** A-B, E) 200 µm; C-D) 50 µm.

**Figure 5.**
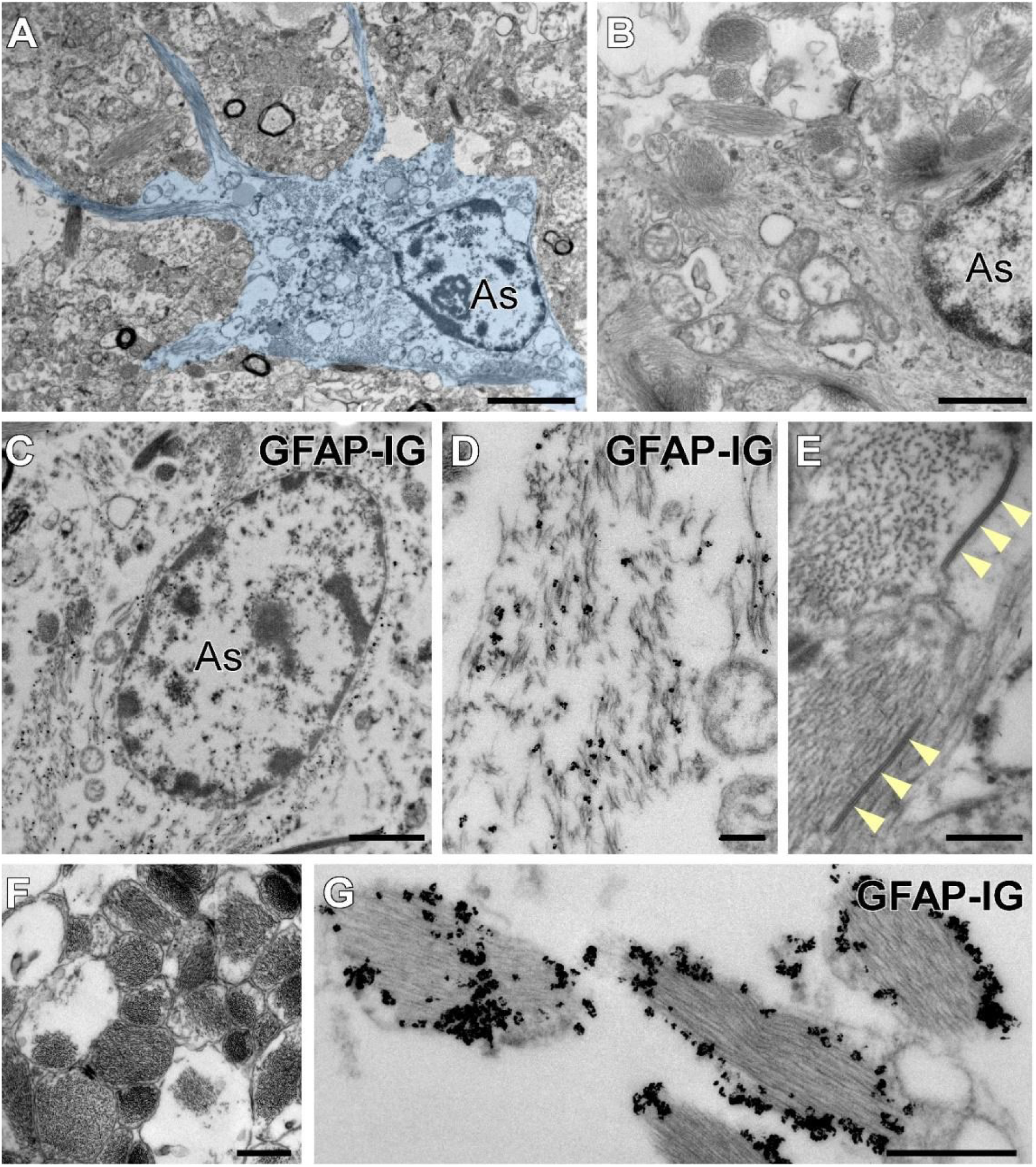
Astrocytic ultrastructure, in the olfactory peduncle (OP). **A)** Ultrastructure of a typical OP astrocyte, pseudocolored in blue, displaying characteristic stellate morphology. **B)** Astrocytes in this region contain dense bundles of intermediate filaments. **C)** Immunogold detection of GFAP confirmed the astroglial nature of this cell population. **D)** Gold particles typically labeled intermediate filaments in the astrocytic cytoplasm. **E)** OP astrocytes frequently formed gap junctions between each other (arrows). **F)** Astrocytic processes in the external layer (EL) exhibited an extraordinarily high density of fibrous components. **G)** These astrocytic processes were distinctly labeled after GFAP immunodetection. The high density of intermediate filaments prevented gold particles from penetrating the tightly packed bundles. **Abbreviations:** As, astrocyte; GFAP-IG, GFAP immunogold detection for transmission electron microscopy. **Scale bars:** A, B) 2 µm; C, D) 1 µm; E, F) 200 nm; G, H) 500 nm.

**Figure 6.**
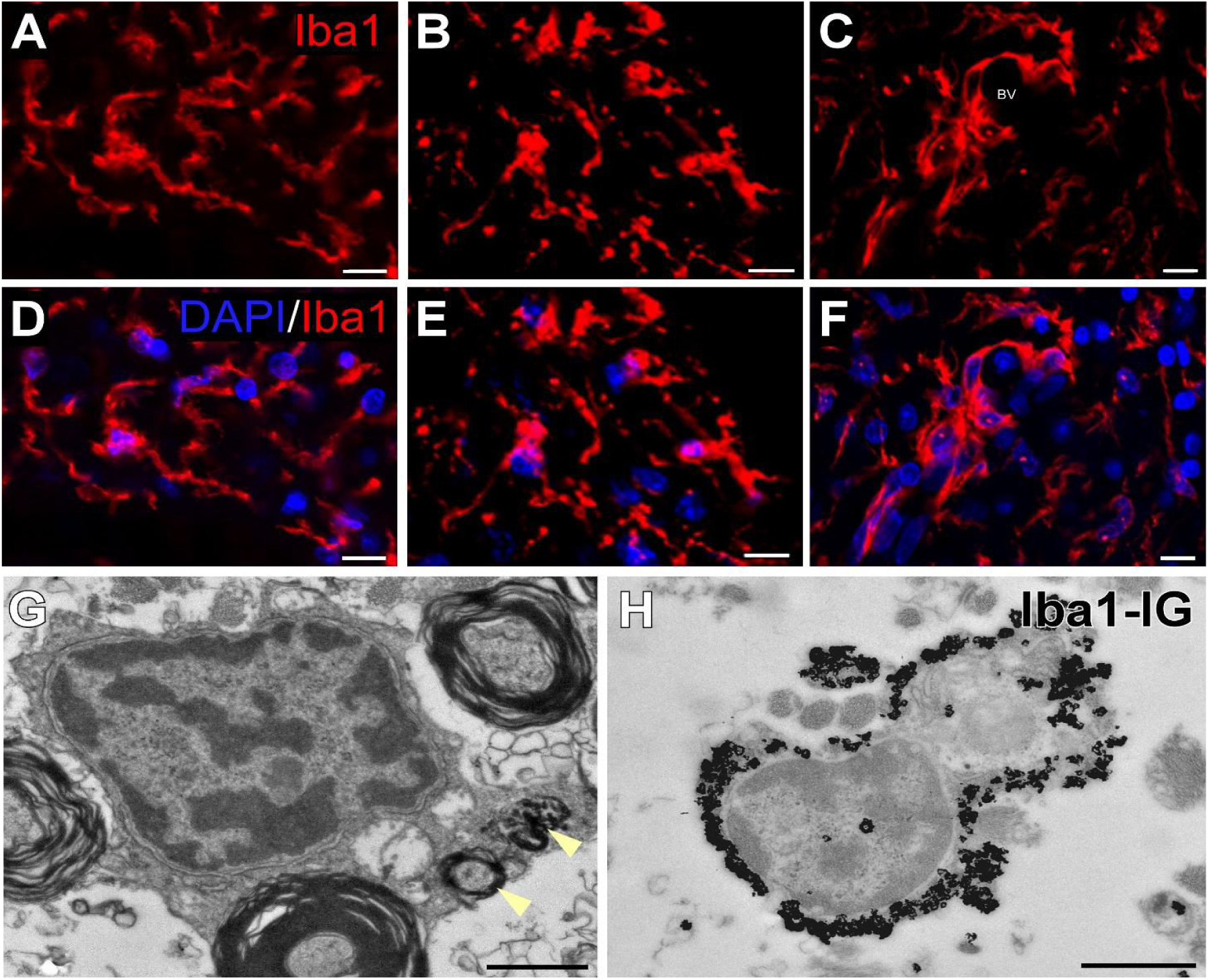
Microglial population in the olfactory peduncle (OP). The OP is populated by microglia, with fewer cells present in the internal layer (IL). Various morphologies of microglial cells have been identified throughout the different layers of the OP. **A, D)** The main morphological type is the resting type, ramified with slender extrusions. **B, E)** Activated morphology: larger, foamy amoeboid cells with reduced extrusions, more frequent in external layers**. C, F)** There are typically microglia accumulations or nodules that can be seen in relation with BVs forming part of the BBB. **G)** Transmission electron microscopy definitively revealed that microglia with typical morphological features, such as lysosomes (arrowheads), were spread across the OP. **H)** Immunogold against Iba1 confirmed the identity of these cells. **Abbreviations:** BBB, brain blood barrier; BV, blood vessel; IG, immuno-gold detection for transmission electron microscopy. **Scale bars:** 5 µm; (J-K) 1 µm.

#### 3.2.2. Axonal layer (AL; Intermediate)

This layer is located between the outer (EL) and inner (IL) layers (Fig. 1C, F) and contains the majority of myelinated axons that comprise the olfactory tract as showed by MBP marker (myelin basic protein) in Fig. 4E. The mean area of this layer in the analyzed sections was 1.01 ± 0.09 mm² (Fig. 2D). This layer contributes approximately half of the total coronal area and occupies the majority of the three layers. The AL area and the fraction that this represents of the total coronal area maintain similar values along the three longitudinal portions studied.

Oligodendrocytes and some astrocytes are present in this layer in association with the myelinated axons. AL is typically annular in shape and surrounds the inner layer tissue (IL). This arrangement is clearly visible in the intermediate portion (Mp) of the OP, although it becomes less distinct as we approach the longitudinal tips (OB or olfactory trigone).

In terms of vascularization, AL showed the highest surface area, as well as the highest number of BVs, with an average count that is more than double that of the other two layers (Fig. 9C). No variation was observed between the longitudinal portions (Fig. 9D). With regard to the BV lumen area (Fig. 9E), no significant variation in the internal area was observed between layers or along the longitudinal axis. The mean values are comparable to those observed in EL or IL (Fig. 9F). The mean percentage of tissue occupied by BV (Fig. 9A) was comparable to the values observed in EL and significantly lower than in IL (Fig. 9G). The density of BV along the OP was consistent (Fig. 9H). As with BVs, a higher count of CA was observed in the AL (Fig. 11C). However, this layer represents the area of the OP that exhibits the lower percentage of CA presence per surface unit (Fig. 11A, G).

#### 3.2.3. Internal layer (IL)

Analysis revealed that the average area for this layer was 0.49±0.06 mm² (Fig. 2D). On average, this represents approximately 25% of the total coronal area. It is the innermost layer, typically enveloped by the AL along the longitudinal axis of the OP. However, this is not universal, as the annular organization of the axonal tract gradually diminishes toward the extremities, particularly in the OB and most notably in the olfactory trigone. It has a lower density of cell nuclei and is mainly composed of numerous dendritic and axonal processes. We observed a lesser presence of astrocytes and only a few isolated myelinated axons, in contrast to what we saw in EL. In this layer, large groups of widely spaced cells with large nuclei are indisputably observed in coronal sections, corresponding to portions of the anterior olfactory nucleus (AON). (Fig. 3G-L). This structure can occupy a significant portion of the coronal section of the OP displacing the other layers.

The vascularization data demonstrate that this layer has a significantly higher BV density compared to the other two layers (Fig. 9A, G). However, it also shows an average BV count comparable to that observed in the EL and significantly lower than that of the AL, as previously stated (Fig. 9C). Furthermore, the mean BV lumen is strikingly similar to that of the EL and AL (Fig. 9E).

This layer also exhibits the highest density of CA (Fig. 11A, G). These are often observed grouped or forming columns that preferentially surround BVs (Figs. 1F, 8E).

### 3.3. Cellular and structural components

In the preceding section, we provided a description of the layers of the OP, together with an account of their characteristics and the principal components that comprise them. Nevertheless, in this section we present data on each of the cellular and structural components that we have identified, noting some of their main characteristics regarding their distribution, relationship with the rest of the structures, and some of their ultrastructural features.

#### 3.3.1. Neurons

For neuron identification, we employed the widely used marker NeuN. Positive cells were primarily located in the AL, with dense accumulations likely associated with the presence of the anterior olfactory nucleus (AON) (Figs. 3G-L, 4A), when present. Under electron microscopy, neurons exhibit abundant organelles, long rough endoplasmic reticulum cisterns, dictyosomes and mitochondria. Neuronal lipofuscin accumulations are frequently observed in these adult human samples (Fig. 3M). The nucleus is typically oval with occasional invaginations, with abundant euchromatin, but also containing small clumps of heterochromatin, often associated with the inner nuclear membrane (Fig. 3N). Axo-somatic synaptic contacts are occasionally visible (Fig. 3O).

#### 3.3.2. Glial cells

##### 3.3.2.1. Astrocytes

Astrocytes, identified by their expression of GFAP (Figs. 4A, C; 5C, D, G), are a highly abundant population found throughout all three layers of the OP. They are uniformly distributed and typically appear as isolated cells, with only a small number of them found in small groups of cells which correspond to the AON. They exhibit a star-shaped morphology, but those located near the surface display a more radial appearance, with one or several radial extensions. Astrocytes in layers AL and IL extend and make contact with BVs. Our study of six adult OP (aged 37-84 years) revealed no glial scaffolds of astrocytes (gliotubes) frequently associated with the rostral migratory stream (RMS), in contrast to what is observed in other adult mammals or during human development and early childhood. TEM analysis showed that astrocytes have an irregular contour and occasional gap junctions with other astrocytes (Fig. 5E). Those in the EL exhibit a more developed cytoplasm compared to those in the AL and IL layers. Additionally, they display intermediate filaments organized into long bundles. These cells contain a few RER cisternae, small dictyosomes, and occasional lysosomes.The nucleus is oval but sometimes irregular, showing invaginations and clumps of heterochromatin (Figs. 5A, C). A frequent feature observed in this postmortem material is the partial absence of the outer nuclear membrane (Fig. 5B, C) probably due to partial degradation prior to fixation.

##### 3.3.2.2. Microglia

We identified microglia with an IBA1 antibody and found them to be present throughout all three portions of the OP, with fewer in the intermediate layer AL. We have identified the three main morphologies (Fig. 6A-F): the “resting” type, which is mainly present and characterized by a ramified shape with slender extrusions; the “activated” morphology, which is larger and foamy with reduced extrusions and is more frequently observed in the external layers; and occasionally, “microglia nodules,” which are accumulations of microglia cells (Schwabenland et al. 2021) that have been usually found associated with BV. The ultrastructure of microglia is characterized by an electron-dense cytoplasm with lysosomes and a nucleus with abundant heterochromatin (Fig. 5G, H).

##### 3.3.2.3. Oligodendrocytes

Oligodendrocyte cells are mainly present in the AL. Using anti Olig-2 antibodies, we observed that these cells occur in a variety of configurations, including as solitary cells, in pairs, and in small groups. (Fig. 7A, B). When studied by TEM, oligodendrocytes appear as small cells in the proximity of myelinated axons. The nucleus is oval and regular in shape, with long clumps of heterochromatin associated with the inner nuclear membrane (Fig. 7C). Their cytoplasm contains some small, dilated cisterns of rough endoplasmic reticulum and few mitochondria (Fig 7D).

**Figure 7.**
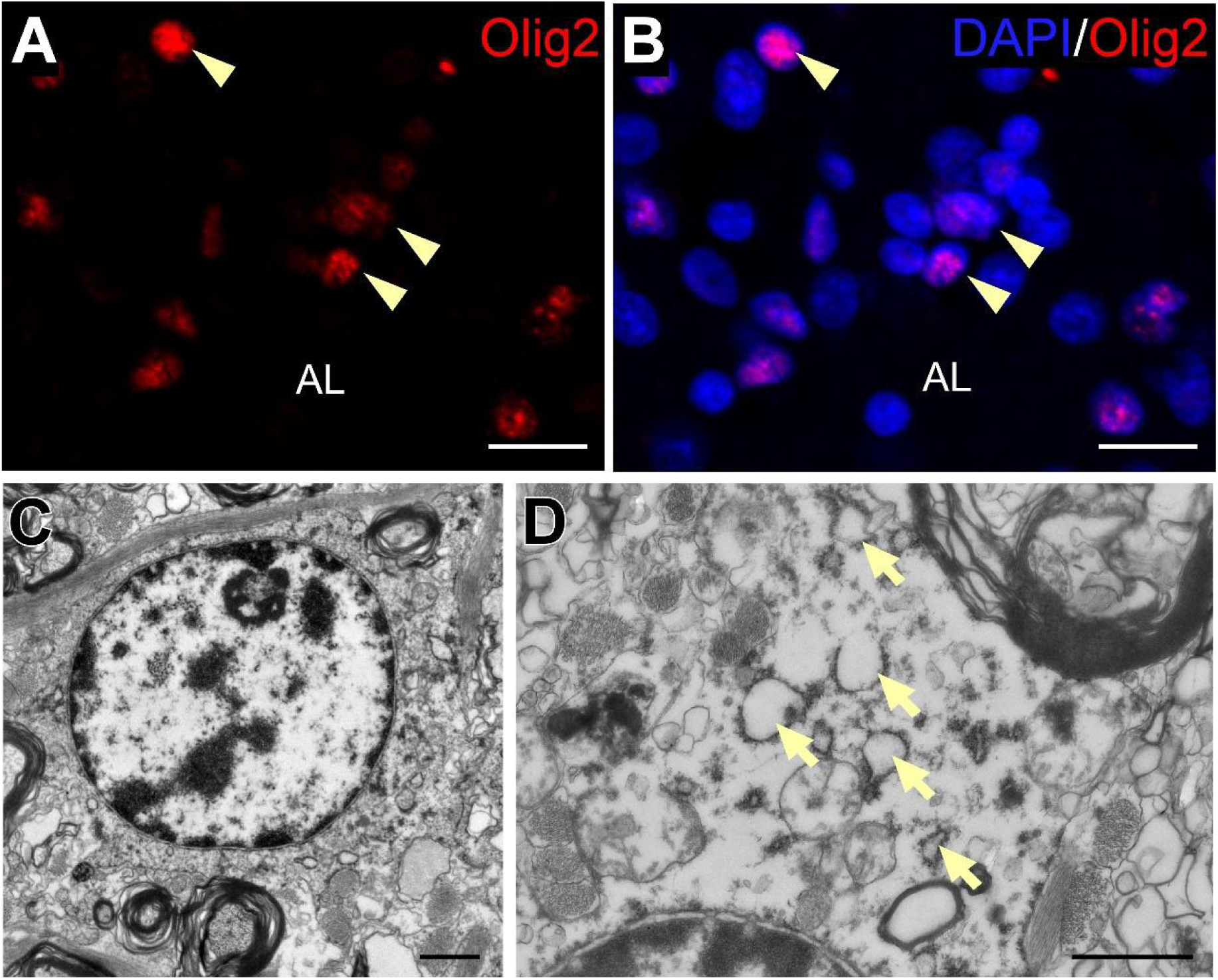
Presence of oligodendrocytes in the olfactory peduncle (OP). **A,B)** Confocal images of Olig2+ cells (arrowheads) in the axonal layer. **C)** A representative oligodendrocyte observed via transmission electron microscopy in the olfactory peduncle. This cell type exhibited a round nucleus with clumped heterochromatin, which consistently covered the inner nuclear envelope. **D)** High-power magnification of an oligodendrocyte cytoplasm, with the presence of typical short and dilated endoplasmic reticulum (arrows). The intimate association with adjacent myelinic axons can also be observed. **Abbreviations:** AL, Axonal layer. **Scale bars:** A, B) 15 µm; C) 1 µm; D) 1 µm.

#### 3.3.3. Blood Vessels

##### 3.3.3.1. Qualitative description

In our observations made using optical microscopy in the analyzed sections of the OP, a large presence of BVs was identified, primarily structures corresponding to capillaries. However, in most of the studied OP sections, three or more larger vessels can also be observed, with diameters close to 10 µm, which, based on their wall structure and size, correspond to arterioles, with a greater presence of smooth muscle, or venules, with fewer muscle fibers but more evident endothelium and fibrous tissue. These larger vessels are more frequent in the EL and IL, while the intermediate AL is richly vascularized, mainly by capillaries. It is very common for these larger BVs to be surrounded by CA.

Using confocal microscopy and specific astrocytic markers (GFAP+), we observed in most vessels astrocytic projections that integrate with the vessel wall, corresponding to perivascular end-feet involved in the blood brain barrier (BBB).

Our TEM analysis revealed that, in addition to typical endothelial cells, OP BVs were associated to multiple astrocytic processes, giving rise to an extensive and intricate network of basal laminae (Fig. 8D). Numerous bundles of intermediate filaments are observed resting on the dense plates. Occasionally, collagen fibers and smooth muscle cells can also be seen as part of the blood-brain barrier (Fig. 8D).

**Figure 8.**
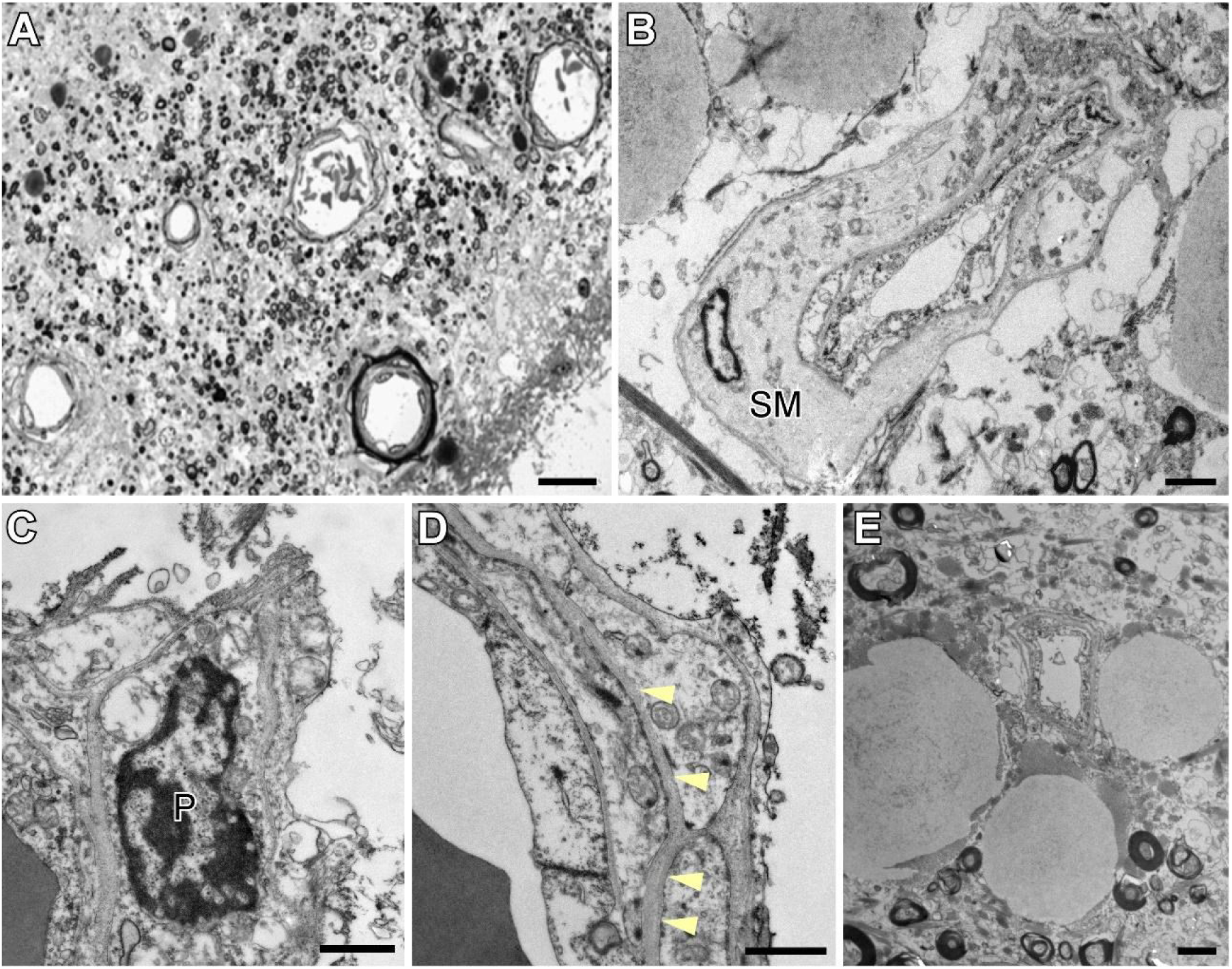
Vasculature characterization in the olfactory peduncle (OP). **A)** Toluidine blue staining of the OP shows that large blood vessels (BV) are located in the peripheral and central zones of the region, predominantly in the external layer (EL) and internal layer (IL). B) Transmission electron microscopy revealed that the majority of BV within the OP exhibited thick walls, characterized by the presence of abundant smooth muscle. **C)** Pericytes (P) were frequently observed. **D)** The presence of multiple layers of the basal membrane is indicative of a tight blood-brain barrier. **E)** The basal membrane was in contact with astrocytes, which were anchored by tight junctions and exhibited a high density of intermediate filaments. Corpora amylacea were observed to accumulate in proximity to BVs. **Abbreviations:** P, pericyte; SM, smooth muscle. **Scale bars:** A) 20 µm; B) 2 µm; C, D) 1 µm; E) 2 µm.

##### 3.3.3.2. Quantitative description

By staining sections with toluidine blue and using image analysis, we were able to quantify the frequency (F_BV_), area (A_BV_), and density (D_BV_) of the BVs identified in 18 coronal sections of six cases in the OP. This analysis was performed for each of the three coronal layers (EL, AL, and IL) and for each of the three longitudinal portions (Ap, Mp, and Pp). For ease of understanding, the statistical results of BV obtained in the different sections, layers and longitudinal portions of OP, have been divided into the following sections: 1) overall count, internal area and density; 2) count by layer and portion; 3) internal area by layer and portion; 4) density by layer and portion.

###### 3.3.3.2.1. Overall blood vessels count, internal area and density

We identified and delimited the luminal areas of all structures compatible with BVs in 18 coronal OP sections, categorizing them into one of the three OP layers: external (EL), axonal (AL), and internal (IL); and one of the longitudinal portions: anterior (Ap), intermediate (Mp), and posterior (Pp).The total number of BVs, represented by F_BV_(X_total_), identified in all sections was 4777. Distribution as follows: EL, F_BV_(X_EL_)= 1098; AL, F_BV_(X_AL_)=2220; IL, F_BV_(X_IL_)= 1459. By longitudinal portions: anterior, F_BV_(X_Ap_)=1558; intermediate, F_BV_(X_Mp_)=1584; posterior, F_BV_(X_Pp_)=1635. The average count per section (Fig. 9C) was F_BV_(X̅_section_)= 265.4±12.5 BVs.

**Figure 9.**
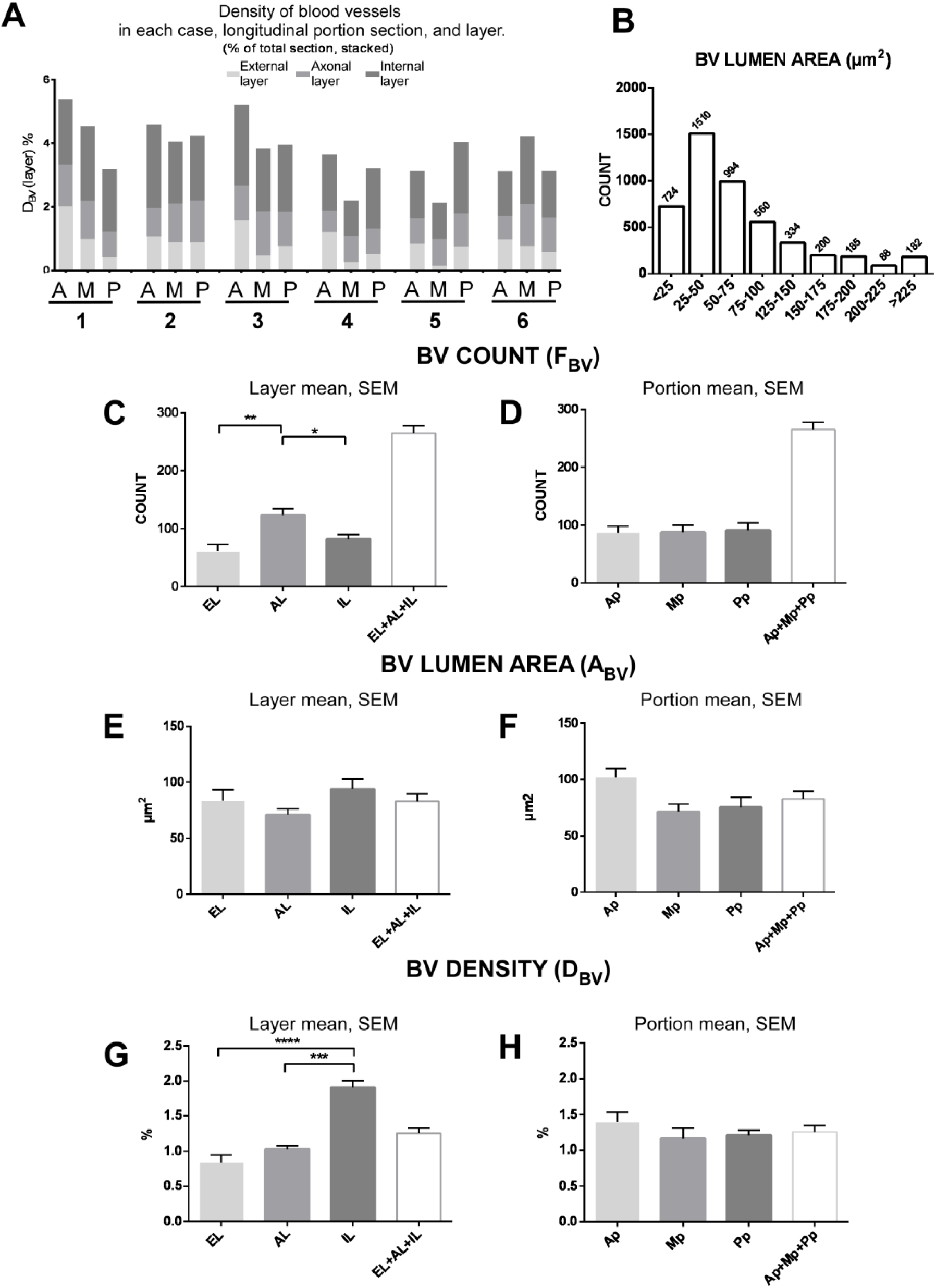
Blood vessel (BV) quantitative description in the olfactory peduncle (OP). For each section, an image analysis was performed to estimate the number (F_BV_), luminal area (A_BV_) and density (D_BV_) of the identified BVs compatible structures. The values obtained have been referenced to the cases, layers, and longitudinal portions in which each element is located. **A)** BVs density, D_BV_, is represented as the percentage of the total surface area occupied by the lumen of the BVs in each layer, longitudinal portion, and case. The mean densities of each layer belonging to the sections of the same case and longitudinal portions have been superimposed. **B)** Luminal area size, A_BV_, of the BVs, frequency histogram. **C)** Average count, F_BV_(X̅_layer_), of the BV identified in each layer and per section (EL+AL+IL). The intermediate axonal layer (AL), which has the largest surface area, exhibits a significantly higher count (KW, P=0.0009; DT, AL vs. EL p<0.001, IL vs. AL, p<0.05) than the other two. **D)** Average count by longitudinal portions, F_BV_(X̅_portion_), in each layer and per section (EL+AL+IL). No differences were found when comparing between sections of the three longitudinal portions. **E)** Comparison of the average luminal area size between the different layers, A_BV_(X̅_layer_), and portions (F), A_BV_(X̅_portion_). The luminal area remains fairly homogeneous, with no significant differences found between the layers or longitudinal portions. Th small differences recorded will affect the results obtained for density. **G, H)** The density, D_BV_, as a proportion of tissue relative to the total size of the layer or section occupied by the BVs lumen. It is shown comparing **(G)** between layers, D_BV_(X̅_layer_), and **(H)** longitudinal OP portions, D_BV_(X̅_portion_). It is observed that the IL layer exhibits a higher density. Although the count for IL has been lower than that obtained for AL and very similar to that of EL (C), the homogeneous size of the BVs lumen, and the smaller surface area presented by this, result in a significantly higher average density value for IL (KW, p<0.0001; DT, IL vs. EL, p<0.0001; IL vs. AL, p<0.001). **Abbreviations:** EL, external layer; AL, axonal layer; IL, internal layer; Ap, anterior portion; Mp, intermediate portion; Pp, posterior portion.

In relation with the lumen of the BVs (A), an average area A_BV_(X̅_section_)= 82.9±6.8 µm^2^ was obtained for BV in the analyzed sections (Fig. 9E). With the data of the recorded individual area values, a frequency distribution histogram has been constructed regarding the measured luminal area for each of the identified BV (Fig. 9B).

The density (D_BV_) of BVs in the OP was evaluated by determining the percentage of coronal area occupied by them, expressed as a percentage of the total analyzed section (Fig 9G). On average, BVs lumen occupied, D_BV_(X̅_section_)= 1.26±0.07 % of the total OP coronal section.

###### 3.3.3.2.2. Blood vessels count by layer and portion

A count has been made of the BVs identified in each tissue section. The obtained values can be associated with each of the cases, the delineated layers, and the longitudinal portions in which they were found. Regarding the average section count of BVs for each layer (Fig. 9A, C): e*xternal layer*, F_BV_(X̅_EL_)= 61.0±11.9*; axonal layer,* F_BV_(X̅_AL_)= 123.3±10.7 BVs; and *internal layer,* F_BV_(X̅_IL_)= 81.1±8.6 BVs. Regarding the average section count of BVs for each portion, data collected suggest that there is no significant BV variation along the OP; not statistically differences in count (F_BV_) have been found along the OP when comparing Ap, Mp and Pp sections.

In relation to the average for layer count in longitudinal portions (Fig. 9D): *anterior portion,* F_BV_(X̅_Ap_)= 86.6±11.9 BVs*; intermediate portion*, F_BV_(X̅_Mp_)= 88.0±11.9BVs; *posterior portion,* F_BV_(X̅_Pp_)= 90.8±12.9BVs.

###### 3.3.3.2.3. Blood vessels luminal area by layer and portion

By tissue section, we have derived the average luminal area of BVs per complete section (Fig. 9E), denoted as A_BV_(X̅_section_), and for each specific layer: A_BV_ (X̅_EL_) for the external layer, A_BV_(X̅_AL_) for the axonal layer, and A_BV_(X̅_IL_) for the internal layer. The gathered data indicate homogeneity in terms of average luminal size, with no significant discrepancies observed upon comparison between the layers (Fig. 9E).

Results concerning the average luminal area by section (A_BV_) of BVs for each of the layers was (Fig. 9E): e*xternal layer*, A_BV_ (X̅_EL_)= 83.7±9.6 µm^2^*; axonal layer,* A_BV_(X̅_AL_)= 71.0±5.3; and *internal layer,* A_BV_(X̅_IL_)= 94.1±9.0 µm^2^. When comparing the average luminal areas of BVs located in each of the three layers, no significant differences have been found (Fig. 9E).

With respect data by portions, no significant differences have been found in terms of luminal area (A_BV_) of the BVs, either in relation to the layer (EL, AL, and IL) or the longitudinal portion (Ap, Mp, and Pp) in which they are located. In case of the average section lumen (A_BV_) of BVs for each longitudinal portions (Ap,Mp,Pp) (Fig. 9F): *anterior portion*, A_BV_ (X̅_Ap_)= 101.9±7.7 µm^2^*; intermediate portion,* A_BV_(X̅_Mp_)= 71.5± 6.7 µm^2^; and *posterior portion,* A_BV_(X̅_Pp_)= 75.4±9.1 µm^2^. Just as the anterior portion shows elevated values, similarly, when comparing layers, no significant statistical differences in luminal size have been established when comparing the longitudinal portions.

###### 3.3.3.2.4. Blood Vessels density by layer and portion

In terms of density (D_BV_), as for the percentage of surface area occupied by the luminal area of BVs, a higher density has been observed in the inner layer (IL). No differences have been found when comparing between layers or longitudinal portions. Regarding the average density (D_BV_) of BVs for each layer (Fig. 9G) was: e*xternal layer*, D_BV_ (X̅_EL_)= 0.84± 0.11%*; axonal layer,* D_BV_(X̅_AL_)= 1.03±0.05%; and *internal layer,* D_BV_(X̅_IL_)= 1.91± 0.10%. Higher density of BVs has been observed in the inner layer (IL) (p< 0.001) (Fig. 9G). As previously mentioned, the mean count of BVs, in the IL does not statistically differ from that of the EL and is even lower than that of AL (Fig. 9C). However, despite the luminal area of the BVs appearing fairly consistent, those in the innermost layer (IL) have a slightly larger internal area, though not statistically significant (Fig. 9E), compared to the other two layers, which may contribute to the increased density observed in the IL (Fig. 9G).

Concerning longitudinal portions (Ap,Mp,Pp) the average density (D_BV_) of BVs for each of them (Fig 9H): anterior portion, D_BV_ (X̅_Ap_)= 1.39±0.14%*; intermediate portion,* D_BV_(X̅_Mp_)= 1.17±0.15 µm^2^; and *posterior portion,* D_BV_(X̅_Pp_)= 1.21±0.14 µm^2^. When comparing the average density among the section groups belonging to each of the three longitudinal portions, we divided the OP into, no significant differences were found (Fig. 9H).

#### 3.3.4. Corpora amylacea

##### 3.3.4.1. Qualitative description

One of the most remarkable features observed in human OP tissue is the presence of CA, visualized as orthochromatic spherical structures with a discernible border and focal areas of increased intensity towards the inner region (Fig. 1D, F). CA are usually found either at the edge of the OP, near the surrounding meningeal tissue (pial/arachnoid), in what appear to be cavities (Fig. 1D), or grouped together, mostly in proximity to BVs (Figs. 1F, 8E), where they form rows that seem to lead toward the cups. CA tend to form groups or rows and are rarely found isolated (Figs. 8E, 1D, F). In some places, these CA accumulations appear to be compressing BVs or axons, which could potentially affect their functionality. In the stroma frequently, CA are surrounded by astrocytic fibers, a finding which has been confirmed by GFAP immunostaining (Fig. 10B, C), and by TEM with anti-GFAP immuno-gold technique (Fig. 10F). In these images, CA can be seen with a spherical morphology and fibrous content, anchored and surrounded by intermediate filaments.

**Figure 10.**
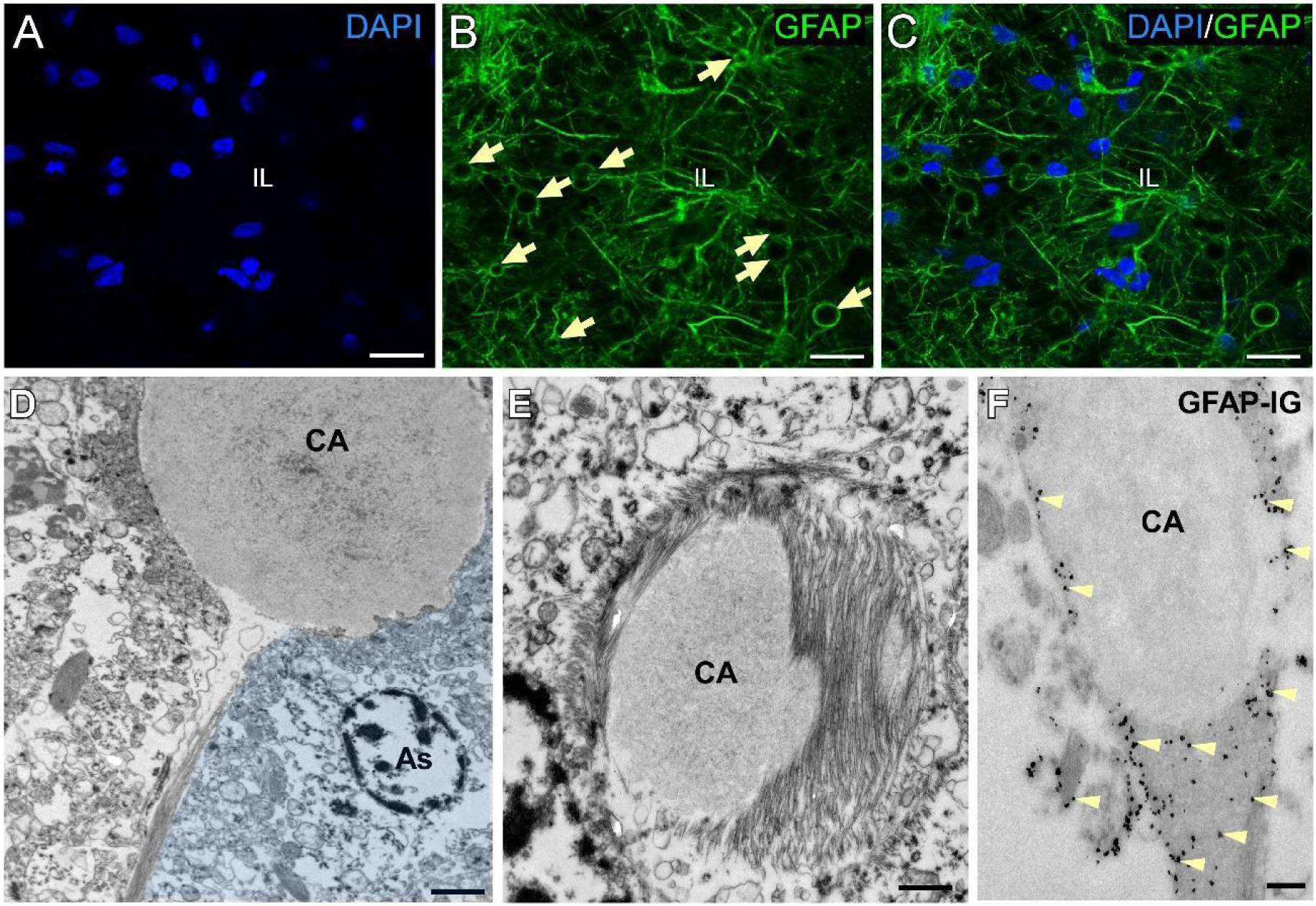
Presence of corpora amylacea (CA) in the olfactory peduncle (OP). **A-C)** Confocal immunofluorescence. Corpora amylacea (CA) are anucleate structures that are typically spherical in shape (arrows) and are surrounded by astrocytic projections (GFAP+). Such structures are often observed in the external layer (EL) of the coronal section, in close proximity to the tissue edge and in the vicinity of blood vessels. **D-F)** Transmission electron microscopy. **D)** CA typically exhibit a racket-shaped morphology, which corresponds to the tip of an astrocytic fibrous process. **E)** At the ultrastructural level, CA exhibit an electron-lucent core surrounded by dense bundles of intermediate filaments. **F)** The intermediate filaments surrounding CA exhibited immunoreactivity for GFAP. Abbreviations: As, astrocyte; CA, corpora amylacea; GFAP-IG, GFAP immunogold detection for transmission electron microscopy. **Scale bars:** A-C) 10 µm; D) 2 µm; E-F) 500 nm.

##### 3.3.4.2. Quantitative description

By staining sections with toluidine blue and using image analysis, as previously discussed for the BVs, we were able to quantify the frequency (F_CA_), area (A_CA_), and density (D_CA_) of the CA identified in 18 coronal sections of six cases in the OP. This analysis was performed for each of the three coronal layers (EL, AL, and IL) and for each of the three longitudinal portions (Ap, Mp, and Pp). For ease of understanding, the statistical results of CA obtained in the different sections, layers and longitudinal portions of OP, have been divided into the following sections: 1) overall count, internal area and density; 2) count by layer and portion; 3) internal area by layer and portion; 4) density by layer and portion; 5) size and density by case.

###### 3.3.4.2.1. Overall *corpora amylacea* count, area and density

The total number of structures identified as CA (F_CA_) in all sections (n=18) was: F_CA_(n)= 18188, is distributed among the 3 layers (EL, AL, IL) with frequencies of: *external layer* F_CA_(X_EL_)= 5843 CA; *axonal layer* F_CA_(X_AL_)= 6094 CA; *internal layer* F_CA_(X_IL_)= 6251 CA. In case of longitudinal portions sections classification: *anterior portion* (n=6, F_CA_(X_Ap_)= 3410 CA); *intermediate portion* (n_portion_=6, F_BV_(X_Mp_)= 8666 CA); *posterior portion*, (n=6, F_BV_(X_Pp_)= 6112 CA). No statistically significant differences were found in the CA count (F_CA_) among the three coronal layers, EL, AL, and IL (Fig. 11C), but significant differences (p<0.01) were observed in the counts referring to the longitudinal portions, Ap, Mp, and Pp (X̅with the anterior portion, Ap, showing the least CA, and Mp sections the higher (Fig. 11D).

**Figure 11.**
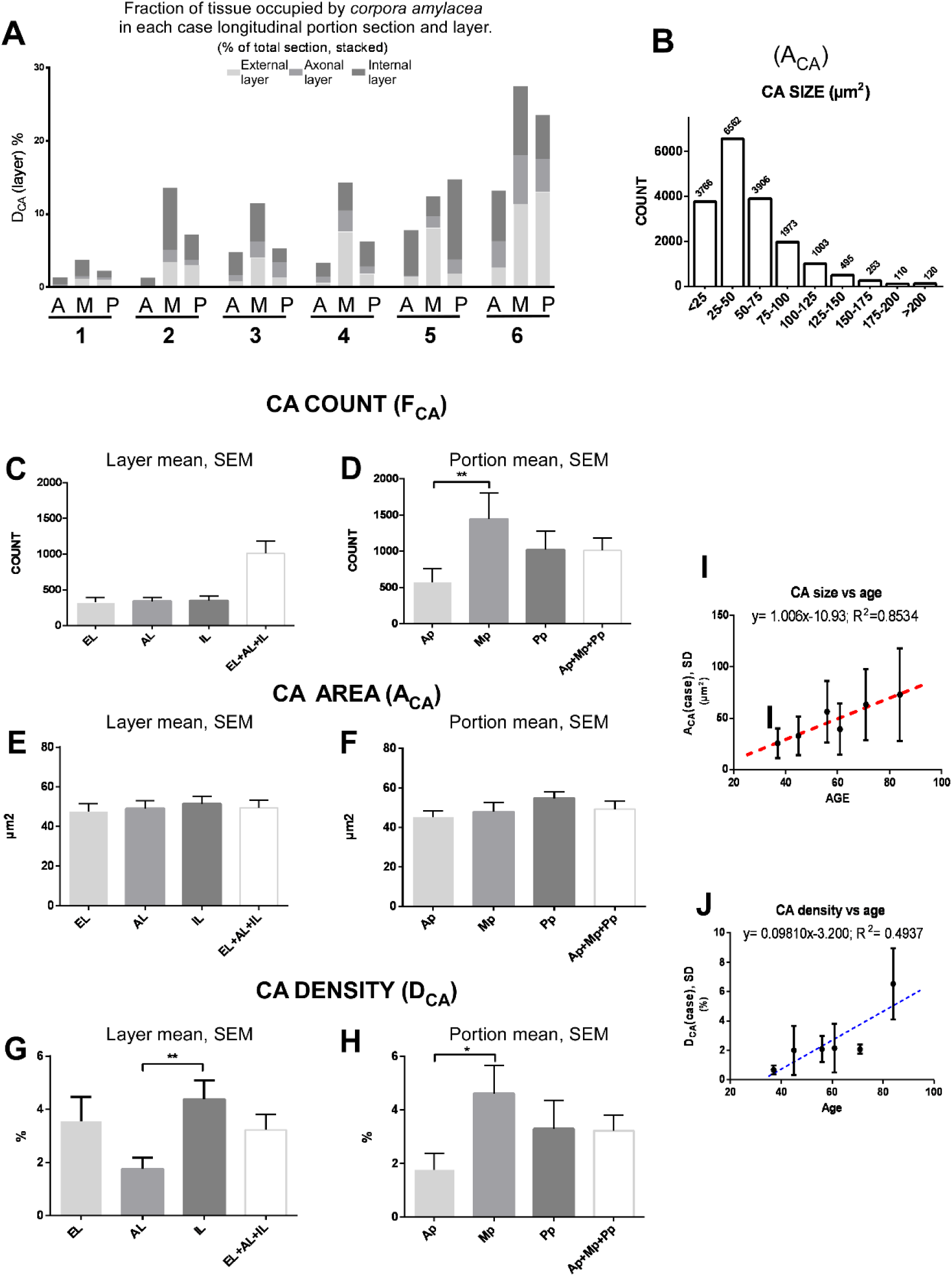
Corpora amylacea (CA) quantitative description in the olfactory peduncle (OP). For each section, an image analysis was performed to estimate the number (F_CA_), size area (A_CA_) and density (D_CA_) of the identified CA compatible structures. The values obtained have been referenced to the specific cases, layers, and longitudinal portions in which each element is located. **A)** The cases under examination have demonstrated variations in the incidence of CA. The density of CA, represented by D_CA_, is expressed as a percentage of the total surface area occupied by them in each layer, longitudinal portion, and case. The mean densities of each layer belonging to the sections of the same case and longitudinal portions have been superimposed for illustrative purposes. **B)** CA histogram of the area size (A_CA_). **C)** No significant differences were observed in the mean count of the CA identified in each layer, F_CA_(X̅_layer_). **D)** The average count by longitudinal portions, F_CA_(X̅_portion_), in each layer and per section (EL+AL+IL) is presented. A comparison of the three longitudinal portions reveals an increased frequency of CA in the intermediate portion (Mp). **E)** A comparison of the average individual area size between the different layers A_CA_(X̅_layer_) and portions **(F)** A_CA_(X̅_portion_) is presented. A comparison of the averages for different layers or sections revealed no differences in size. A similar observation was made when comparing between longitudinal portions. **G, H)** The density, D_BV_, is expressed as a proportion of tissue relative to the total size of the layer or section occupied by the CA. The data in **(G)** is presented for comparison between layers, D_CA_(X̅_layer_), and **(H)** for longitudinal portions, D_CA_(X̅_portion_). It is observed that the IL layer exhibits a higher density despite the CA count or size for IL being very similar to that obtained for the others. The intermediate portion has demonstrated a markedly elevated density (H) and frequency (D) of CA. **I, J)** Linear regressions on the variable age in relation to size and density. There is a strong correlation between these variables, but not with frequency. We can observe that the size of the CA shows a better fit, indicating a clear increase in size with advancing age of the individuals. **Abbreviations:** EL, external layer; AL, axonal layer; IL, internal layer; Ap, anterior portion; Mp, intermediate portion; Pp, posterior portion.

The average CA count per section was F_BV_(X̅_section_)= 1010± 173 CA (Fig. 11C, D). In regard to the individual area of the CA (A), an average area of A_CA_(X̅_section_)= 49.31±3.93 µm^2^ was obtained for CA in the analyzed sections (Fig. 11E, F). For individual CA sizes, a frequency distribution histogram has been created (Fig. 11B).

Concerning density (D_CA_), we estimated that the CA occupy an average of 3.23±0.58% of the total coronal section of the OP (Fig. 11A, G, H). Although there are clear differences in CA counts between sections and cases, a statistically significant relationship between the count of CA and the age of the subjects could not be established.

###### 3.3.4.2.2. *Corpora amylacea* count by layer and portion

The average section counts of CA for each layer (Fig. 11C) was: e*xternal layer*, F_CA_(X̅_EL_)= 325±71*; axonal layer,* F_CA_(X̅_AL_)= 339±56 CA; and *internal layer,* F_CA_(X̅_IL_)= 347±66 CA. The average section counts of CA for each longitudinal portion (Fig. 11D) was: anterior portion, F_CA_ (X̅_Ap_)= 568.3±188.9*; intermediate portion,* F_CA_(X̅_Mp_)= 1444.0±357.7; posterior portion, F_CA_(X̅_Pp_)= 1019.0±256.1 µm^2^.

###### 3.3.4.2.3. *Corpora amylacea* area by layer and portion

CA show an average size of A_CA_(X̅)= 49.31±3.93 µm2 (Fig.11E, F) without significant changes when comparing their size between layers or longitudinal portions. Assuming a completely spherical morphology of the CA, something that is not entirely accurate, this individual size value would correspond to an approximate average diameter of, CA_Ø_= 8 µm.

The average CA individual area (A_CA_) by section of CA for each layer (Fig. 11C) was: e*xternal layer*, A_CA_ (X̅_EL_)= 47.40±4.11 µm^2^*; axonal layer,* A_CA_(X̅_AL_)= 49.06± 3.94 µm^2^; and *internal layer,* A_CA_(X̅_IL_)= 51.36± 3.892 µm^2^. No statistically significant differences were found among the average sizes when comparing the 3 layers (EL, AL, IL).

Regarding average, individual area (A_CA_) by section of CA for each portion (Fig. 11D) was: *anterior portion*, A_CA_ (X̅_Ap_)= 45.19± 3.30 µm^2^*; intermediate portion,* A_CA_(X̅_Mp_)= 47.84± 4.79 µm^2^; *posterior portion,* A_CA_(X̅_Pp_)= 54.80± 3.36 µm^2^. No statistically significant differences were found among the average CA sizes when comparing the 3 longitudinal portions (Ap, Mp, Pp).

Furthermore, we have observed that the CA average area per clearly increases with the age of the subjects (ranging from 37 to 84 years old) (Table 1A), showing a strong correlation (r= 0.9429, p<0.01). A linear regression line has been recalculated, relating CA area to age (y= 1.006x-10.93; R^2^= 0.8534) (Fig. 11I).

###### 3.3.4.2.4. *Corpora amylacea* density by layer and portion

For each layer, data on average CA density (Fig. 11A, G) per section was: *external layer*, D_CA_ (X̅_EL_)= 3.55± 0.92%; *axonal layer*, D_CA_(X̅_AL_)= 1.76± 0.41%; and *internal layer*, D_CA_(X̅_IL_)= 4.37± 0.72%. This IL exhibits a significantly higher density than that obtained for AL. The statistical analysis revealed significant differences when comparing the average density among the three layers (Fig. 11G), with IL exhibiting a higher density than AL (p<0.01).

Comparing the average CA density per section with respect to the longitudinal portions (Fig. 11A, H), anterior portion, D_CA_ (X̅_Ap_)= 1.764± 0.48 %; intermediate portion, D_CA_(X̅_Mp_)= 4.618± 0.77%; posterior portion, D_CA_(X̅_Pp_)= 3.30± 0.83%), the intermediate portion (Mp) reveals a significant higher proportion than Ap of tissue occupied by these (p<0.05).

###### 3.3.4.2.5. *Corpora amylacea* size and density by case

Based on the data obtained, we decided to employ density as a comparative measure between cases. Similar to the relationship observed between the increase in average CA size, A_CA_(X̅_case_), and age, a correlation was also found between the average density of occupied CA, D_CA_(X̅_case_), and the age of individuals, (r= 0.834; p< 0.05) (Fig. 11J). However, the correlation coefficient and goodness of fit for the regression line (y= 0.09810x-3.200; R^2^= 0.4937) were found to be lower than that obtained when comparing size/age (Fig. 11I).

## 4. Discussion

### 4.1 The anatomical and cellular organization of the olfactory peduncle: implications for its physiology

The olfactory system has been extensively studied in many mammalian species, but in humans, perhaps due to our microsmatic nature or the lack of consensus regarding the identification of human pheromones, it has been remained comparatively less explored (Wyatt 2015). In the study of this system, the OB has classically been the olfactory structure that has received the most work, as it is the first relay station for olfactory information from the primary olfactory neurons in the olfactory epithelium. It is important to highlight that in recent years, numerous studies have also concentrated on findings related to OB plasticity and its potential as an adult human neurogenic niche (Maurice A. Curtis, Kam, y Faull 2011; Huart, Rombaux, y Hummel 2013; Bonfanti et al. 2023).

The olfactory information is transmitted to the OB via the cribriform plate of the ethmoid bone, where it is processed in the glomeruli. This information is transmitted centripetally along the OP, also known as the olfactory tract, to the primary olfactory cortex, which includes brain regions like the piriform and entorhinal cortices. The OP functions as the main conduit for these signals, with further projections from the primary olfactory cortex to regions such as the orbitofrontal cortex and hippocampus. Additionally, the OP carries centrifugal fibers from areas including the contralateral OB, the diagonal band, and other brainstem nuclei(Paxinos 2014). It is therefore essential to have a clear understanding of the established normal characteristics of the OP, as well as any potential anatomical, physiological or histological changes that may occur. Any alterations in this region could potentially lead to pathological conditions or loss of sensitivity, the aetiology of which is not yet fully understood.

In terms of morphology, the OP can be defined as the slender stalk of tissue that connects the olfactory bulb to the ventral forebrain. It plays a pivotal role in primates, as evidenced by its presence in both humans and macaques, where it connects the OB with olfactory cortices. Defined topologically through dissection, the OP lies on the ventral surface of the orbital gyri, nestled within the olfactory sulcus, between the straight gyrus (medially) and the medial orbital gyrus (laterally). The OP contains components of the olfactory tract and the anterior olfactory nucleus, which are integral to the transmission of olfactory signals (Paxinos 2014). With regard to embryonic development, the OP along with the OB, are projections from the prosencephalon. Both structures are covered by the meninges: dura mater, arachnoid mater, and pia mater. The development of these structures commences around the sixth week of gestation, when olfactory axons migrate towards the prosencephalon. By the 14th week, the histogenesis of the OB is underway, with initial layer organization becoming observable. Full neuronal maturation continues until birth and beyond, maturing until approximately the age of 20 years, when they typically reach their definitive size (Sarnat y Flores-Sarnat 2017).

Therefore, the aim of this study is to present information on both macroscopic and especially microscopic characteristics of the OP, based on six individuals, three males and three females, ranging from 37 to 84 years of age (Table 1A), all undiagnosed with neurological pathologies.

With respect to the external morphology of the OP, measurements from the analyzed cases show that the length of this structure varies between individuals and may also exhibit asymmetry within the same individual. Literature references mention such size differences, which are often considered physiologically normal and without impact on olfactory function, possibly due to compensatory mechanisms (Manan et al. 2022). In our study, although control cases were utilized, both undiagnosed with neurological conditions in life and post-mortem (Braak), we cannot determine whether the observed asymmetries (left-right) in the OP were congenital or developed during the lifespan. Moreover, although we have divided the OP into three longitudinal portions (Ap, Mp, and Pp) for differential analysis, these do not precisely correspond to the morphological changes or cytoarchitecture that we could identify. However, it is clear that there is a longitudinal variation in the external coronal morphology of the OP, as well as in the distribution or size of the histological layers, which essentially corresponds to the three types presented (Fig.1B), depending on whether we observe peribulbar (Ap), intermediate (Mp), or retrobulbar (Pp) areas (Crosby y Humphrey 1941) (*supp. mat. 8*). This generally allows identification by appearance of whether a coronal section corresponds to any of these portions. In terms of the arrangement of the layers into which we have divided the OP, this study observed, in the majority of the sections analyzed, a circular arrangement of the olfactory tract that runs through the OP. This arrangement seems to diminish in areas very close to the anterior limits, the olfactory bulb, and posteriorly at the olfactory trigone. We have designated this layer of the olfactory tract as the AL, which delineates a more external region, the EL, and a more internal region, denoted as IL. This classification has allowed for a differential study of the sections of the OP. Common elements among the layers include various cytoarchitectural elements that play a role in the structural and functional support of nerve cells, contributing to the tissue’s homeostasis. This correlates with the observed vascular supply; although large vessels were primarily found in the internal layer, with almost no presence in the AL, capillaries are evident throughout the entire structure.

Pertaining to cellular elements, neurons are primarily located in the AL, with dense accumulations likely associated with the presence of the anterior olfactory nucleus (AON) when present in the IL. Neurons are also frequently observed in the EL and, to a lesser extent, in the IL, where some isolated or small groups of neuronal somas can be found outside the main tract in the AL. Ultra-structurally, lipofuscin accumulations are frequently observed in neurons in these adult human samples. Astrocytes are the most abundant cell population in the OP and are present in all three layers, typically with a stellate morphology. However, their morphology and organization show some variations. In the EL, particularly in areas close to the meningeal tissue surrounding the OP, astrocytes display a more radial morphology, with a more developed cytoplasm and intermediate filaments organized into long bundles, suggesting greater activity and specialization in this more exposed layer. In contrast, those found in the AL have shorter filaments, similar to the few observed in the IL. Regarding other glial elements, microglia are evident and exhibit a homogeneous distribution throughout the OP, whereas oligodendrocytes are restricted to the AL. In terms of cell density across the layers of the OP, the IL shows a lower presence of cells, being primarily composed of numerous dendritic and axonal processes.

From a physiological perspective, the OP undergoes various changes throughout life, one of which is the presence of a RMS. During development, the OP receives migrating neuroblasts that travel tangentially from the ventricular-subventricular zone (LV-SVZ) of the anterior horn of the lateral ventricles toward the OBs, encased in a richly vascularized astrocytic scaffold. These neuroblasts mature, integrating radially into the olfactory circuitry of the OB. In some adult mammals, this migration persists, though to a lesser extent. In human adults, studies suggest that this migration becomes highly limited, virtually ceasing after adolescence. However, it remains a controversial topic whether remnants of these structures persist in the adult human and with what functionality they might exist. (Sanai et al. 2011; 2004; M. A. Curtis et al. 2007). In this regard, in the cases we have evaluated, no structures reminiscent of the astrocytic scaffold that forms the migratory route of the RMS have been found, supporting the idea that this structure does not persist in adult humans.

Regarding physiopathology, some studies correlate dysosmias or specific neuropathologies with atrophies or volume changes, including cases of partial or complete agenesis, but typically focus on the OB (Yousem et al. 1996; Buschhüter et al. 2008). However, scarce data were found linking specific OP cytoarchitecture alterations or minor length variations with pathological conditions. Furthermore, conditions such as hyposmia or anosmia are proposed as indicators of underlying neurological or systemic conditions, including Alzheimer’s disease, Parkinson’s disease, or SARS-CoV-2 infection, but currently lack clear diagnostic potential. Therefore, a comprehensive understanding of the anatomy, functionality, and physiology of the OP is essential for accurate diagnosis and effective treatment of these disorders.

### 4.2. Tissue posfixation

The principal objective of tissue fixation is to maintain the integrity of cells and tissue components in a state that most closely resembles their natural form. This is achieved by preventing autolysis, which commence as soon as a tissue is deprived of its blood supply, thereby facilitating the preparation of thin sections. In our case, once fixed by perfusion, human brain tissue is subjected to a 4% paraformaldehyde postfixation of indeterminate duration for storage. This method has proven valid for ultrastructural studies even after 5 years of postfixation (Insausti et al. 2023). However, we have encountered some difficulties in detecting certain antigens using immunostaining techniques. In this regard, authors such as Engel K. et al. 2011 or (Piña et al. 2022) have reviewed some approaches that improve immunostaining and better antigen recognition with postfixation durations of 30 minutes to 2 hours compared to 24 to 48 hours. This supports the theory that post-fixation time is a limiting factor in achieving good antigen preservation, enhancing antibody sensitivity and facilitating the obtaining of improved signals in immunolabeling techniques. Our findings align with this theory since we have observed that prolonged post-fixation times can result in excessive protein cross-linking, which in turns reduce antigen accessibility and signal intensity. This conclusion is in accordance with (Maraziotis et al. 1992) who described enhanced staining for synaptophysin with 1 hour postfixation versus 24 hours. These studies suggest that shorter postfixation times are generally more effective for some epitopes, though the optimal duration may depend on the tissue and antibodies used, requiring further optimization for specific protocols and brain regions.

These tissues generally are typically derived from cases where varying time periods may have elapsed between death and fixation, as well as differences in the type, conditions, and duration of tissue preservation from the time of collection to experimental use.

### 4.3. Blood perfusion in the human olfactory peduncle

The olfactory epithelium and ventral side of the OB receive crucial extracranial blood supply from the sphenopalatine artery, as well as the anterior and posterior ethmoidal arteries (Bartholdy 1897; Osborn 1978). Intracranially, the olfactory artery attaches to the OP and OB through the arachnoid, thereby supplying the entire structure (Favre et al. 1995; Hendrix et al. 2014). Nevertheless, to the best of our knowledge, no previous studies have provided information about the internal distribution, lumen area, or density of the BVs in the human OP, as we present here. We have observed capillaries in all layers, and in our quantification, BVs appear to have a relatively stable lumen throughout the OP and across the three described coronal layers, with a significant increase in density in the IL, where some larger arterial vessels are present. However, we were unable to obtain clear images of the perforating branches in the stroma OP from these large blood vessels, which could potentially show the origin of some of these capillaries. Additionally, the axonal tract running through the AL is well vascularized. Disruptions in the intrapeduncular blood supply in this region could lead to dysosmias, affecting the conduction pathways due to hypoxia-mediated neuronal apoptosis or inflammatory processes resulting from the penetration of substances into the brain parenchyma. Furthermore, the ultrastructure of the BBB in the olfactory epithelium, OB, and, seemingly, in the OP differs from that observed in the remainder of the central nervous system. A comprehensive review of the existing literature revealed a paucity of detailed descriptions concerning the cytoarchitecture of the human OP. In this regard, we observed the presence of microglial nodules associated with the walls of certain BVs; however, we did not delve into the specific characterization of the BBB ultrastructure or its differences between layers or longitudinal portions. Nevertheless, it is well documented that the blood-brain barrier in this region is more permeable than in other areas of the central nervous system, thereby facilitating the diffusion of external particles or pharmacological agents (Thorne y Frey 2001).

It is important to emphasize that qualitative observation and the quantification system used in this study, which relied on coronal sections of the OP, suggests that the majority of BVs run along the anteroposterior axis of the OP. However, not all follow this orientation, which may affect the appearance of the luminal area in coronal sections, potentially increasing the values when extrapolating the luminal diameter. To improve measurement accuracy, vessels that were clearly sectioned along these planes were excluded from the analysis to avoid skewing the data. In addition, the Nissl stains used for vessel quantification did not allow for the distinction between arterial and venous components. Furthermore, it is possible that some structures quantified by optical microscopy as capillaries may actually correspond to elements of the glymphatic system. During this study, we encountered suboptimal results with immunohistochemical approaches and various anti-laminin antibodies for blood vessel identification in our tissues. In future analyses, the use of specific markers, such as CD31 (Bergom et al. 2008) or CD34 (Naeim 2008), which label endothelial cells of BVs, or podoplanin (D2-40), a marker for identifying lymphatic capillaries (Yokomori et al. 2010), is being evaluated. These markers, along with 3D confocal reconstructions and ultrastructural analysis via electron microscopy, will be crucial for further characterizing the vascular network of the OP. However, given the lack of available information regarding the internal vascularization of the olfactory OP, we believe that the data presented may provide a foundation for more precise future stereological studies.

### 4.4. *Corpora amylacea* in the human OP

*Corpora amylacea* are mainly composed of polyglucans and can be identified in a multitude of organs in adult humans, including the lungs, prostate gland and also throughout the central nervous system, such as the brain, midbrain, cerebellum, and spinal cord. The presence of these structures is linked not only to normal aging but also to various pathophysiological processes. In the brain, CA can be observed in regions such as the hippocampus, substantia nigra, cerebral cortex, and olfactory peduncle, indicating a widespread distribution. They are often found in proximity to ventricular walls, pial surfaces, and BVs (Alder 1953; Sakai et al. 1969b; Sbarbati et al. 1996; Navarro et al. 2018; Riba et al. 2021). An understanding of these locations is essential for grasping the role and clinical significance of CA in normal aging but also in neurological conditions.

Given their high prevalence in the OP (Bathini et al. 2019), this study examines their distribution across the various layers, as well as their size and density. CA have been identified in all sections of the OP in our cohort, including 6 individuals (ranging from 37 to 84 years old) without diagnosed neuropathology. A correlation between age and the proportion of tissue occupied by the CA, but particularly between age and the size of the CA. However, a statistically significant correlation between age and CA count could not be established.

It is noteworthy that in one case excluded from this study due to its condition, a 94-year-old male post-mortem diagnosed with Braak 3 Alzheimeŕs disease (Braak y Braak 1991), both the count and density of CA were markedly low (data not shown), aligning with those observed in the youngest subject (37 years old) included in our cohort. However, the average individual size of the CA did resemble the values presented for these ages. Based on these data, we hypothesize that there may be a relationship between CA and Alzheimeŕs disease or other neurodegenerative pathologies, whether CA are a cause or consequence of neurodegeneration, and whether they are risk or protective factors in neurodegenerative processes.

The aggregation of proteins such as Tau or β-amyloid has been observed in CA, and several authors have proposed a correlation between the number, size, or composition of these proteins and central nervous system diseases. However, the results are insufficient and sometimes contradictory, thus rendering their interpretation in the context of these disorders and their diagnostic utility (Rohn 2015). As a result, their function and clinical relevance remain unclear. There is still limited knowledge about their presence and role in normal physiological conditions (Mrak, Griffin, y Graham 1997). There seems to be a consensus that they accumulate substances, which can also be found in the neuropil, acting as “waste containers” or “wasteosomes” (Riba et al. 2021). However, the mechanism by which molecules are incorporated is also unknown.

Regarding their origin, it has been postulated that CA are formed as a consequence of cellular degradation (Ramsey 1965; Cavanagh 1999; Augé et al. 2018). The presence of organelles and molecules associated with neuronal or axonal degradation (Anzil et al. 1974) has been documented in the literature yet there is no consensus on this matter (Galli et al. 2018). With regard to this aspect, our findings indicate that CA are not present within axons or neuronal somas. Conversely, as previously proposed by other authors and demonstrated in this study, there is a clear relationship between these structures and astrocytes of the OP (Ramsey 1965; Wander et al. 2022). However, this has not always been the case. It is unclear whether this is because they are not always related to the glia and CA can remain independent, or due to histological circumstances that have not allowed for the correct identification. Nevertheless, this study provides strong confirmation that they can be confined by intermediate filaments (GFAP+) from astrocytic projections (MET-G and confocal techniques), which may suggest a potential astrocytic origin (Fig. 10).

Some authors have also proposed that CA may be susceptible to be phagocytosed, by microglia/macrophages (Suzuki et al. 2012). However, this hypothesis has not been confirmed in this study in the OP. In none of the TEM or IF images observed with confocal microscopy and conducted with anti-IBA1 antibodies, a marker of both resting and activated microglia, were CA located in association with these cell types.

Another noteworthy aspect is the distribution of CA in the OP, which is evidently not random. In the central nervous system, the presence CA has typically linked to their proximity to pial tissue, the ventricular system, and BVs (Navarro et al. 2018). This suggests that the tissue in question is capable of creating or displacing the CA to these locations, which can be easily associated with drainage. There has been speculation regarding the possibility that astrocytes are the responsible for forming and we ask if they could be displacing them to these locations through their projections (Fig. 10), as they appear to be closely related to the reorganization of the extracellular matrix (Silver y Miller 2004).

Our observations of OP sections indicate that CA are often located in close proximity to BV. Additionally, they tend to accumulate at tissue boundaries, particularly near the pial tissue that extends into the subarachnoid space, in what appear to be cavities at the edges of the OP (Fig. 1D), which may be organized collagen fibers that play a role in the neuropil-pia-subarachnoid transition to CSF and bloodstream for CA elimination. In this regard, their presence has been detected in cerebrospinal fluid (Cavanagh 1999; Riba, Campo-Sabariz, et al. 2022).

As previously discussed, there is a need to enhance our understanding of these structures and their physiological functions, particularly if correlations with pathologies are established based on their presence. Another aspect deserving investigation is the development of techniques for in vivo characterization. Some researchers are attempting to detect these structures in human brains using non-invasive methods such as magnetic resonance imaging (MRI) (Abel et al. 2010; Pimentel, Roque, y Cunha E Sá 2020) or synchrotron radiation X-rays (Lee et al., 2023).

Therefore, according to the research presented here, there are still many challenges and unanswered questions that require a deeper study of changes in morphology, the cytoarchitectonic components of the OP, and their modifications over time. Such research may better explain their physiological implications in the processes of normal or pathological aging and the potential therapeutic approaches for the various conditions underlying the common partial or total loss of smell that occurs with age.

## Abbreviations

AL: Axonal-intermediate layer of the olfactory peduncle
AON: Anterior olfactory nucleus
Ap: Anterior longitudinal portion of the olfactory peduncle
As: Astrocyte
BBB: Blood brain barrier
BV: Blood vessel
CA: *Corpora amylacea*
EL: External layer of the olfactory peduncle
IG: Immunogold detection for transmission electron microscopy
IL: Internal layer of the olfactory peduncle
LRC: Slices collection solution
LV: Lateral ventricle
Mp: Middle (intermediate) longitudinal portion of the olfactory peduncle
MT: Meningeal tissue
N: Neuron
P: Pericyte
Pp: Posterior longitudinal portion of the olfactory peduncle
RMS: Rostral migratory stream
SM: Smooth muscle
SVZ: Subventricular zone
TEM: Transmission electron microscopy

## 5. Acknowledgments

This work has been funded by the Regional Government of Castilla-La Mancha/European Regional Development Fund (young researchers grant no. SBPLY-19-180501-000216) to CRP and by the Valencian Council for Education, Culture, University and Employment (PROMETEO/2023/053), CIBERNED (CB06/05/1131), ISCIII, MCIN, and European Union – European Regional Development Fund to JMGV.

## 6. Authors’ contributions

JFRG, SGG, CRP and JMGV carried out all the experiments. JFRG, CRP, VHP, JMGV wrote the first draft of the manuscript. EA contributed to data analysis. VHP, ACC and RI contributed to section 1 and all figures as well as to edition and English revision of the manuscript. CRP and JMGV coordinated the work. All authors have read and approved the final manuscript.

---

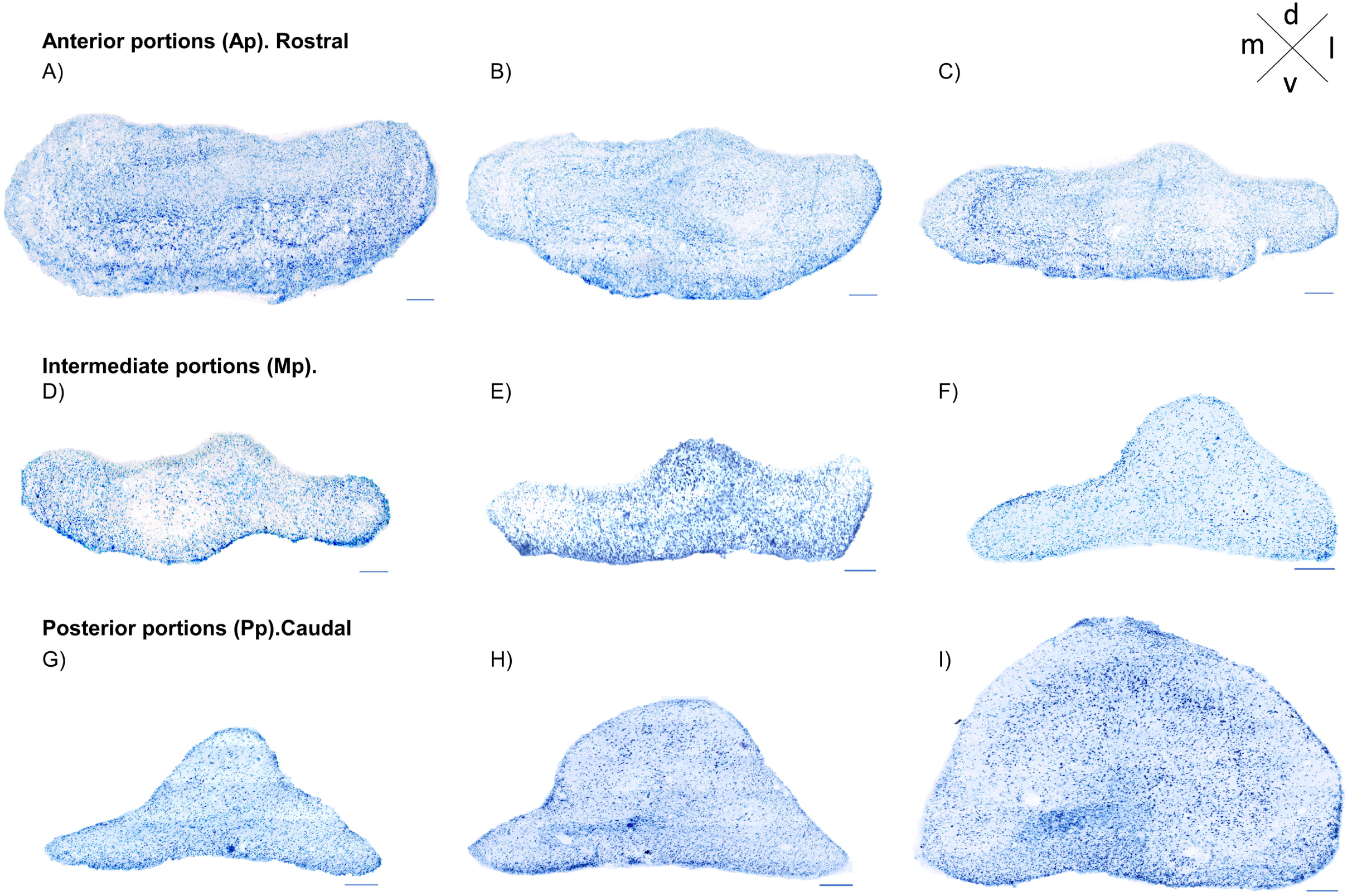
A-1) Human olfactory peduncle, from A) peribulbar regions to the I) rostral part of the olfactory trigone. A-C) Anterior portion sections; D-F) intermediate portion sections; G-1) posterior portion sections. Panoramic 20x, coronal sections; 50µm thickness. Tionine blue. Distance between images, 400 mm. Scale bars: 200µm

